# Transcriptional landscape of human microglia reveals robust gene expression signatures that implicates age, sex and *APOE*-related immunometabolic pathway perturbations

**DOI:** 10.1101/2021.05.13.444000

**Authors:** Tulsi Patel, Troy P. Carnwath, Xue Wang, Mariet Allen, Sarah J. Lincoln, Laura J. Lewis-Tuffin, Zachary S. Quicksall, Shu Lin, Frederick Q. Tutor-New, Charlotte C.G. Ho, Yuhao Min, Kimberly G. Malphrus, Thuy T. Nguyen, Elizabeth Martin, Cesar A. Garcia, Rawan M. Alkharboosh, Sanjeet Grewal, Kaisorn Chaichana, Robert Wharen, Hugo Guerrero-Cazares, Alfredo Quinones-Hinojosa, Nilüfer Ertekin-Taner

## Abstract

Microglia have fundamental roles in health and disease, however effects of age, sex and genetic factors on human microglia have not been fully explored. We applied bulk and single cell approaches to comprehensively characterize human microglia transcriptomes and their associations with age, sex and *APOE*. We identified a novel microglial signature, characterized its expression in bulk tissue and single cell microglia transcriptomes. We discovered microglial co-expression network modules associated with age, sex and *APOE*-ε4 that are enriched for lipid and carbohydrate metabolism genes. Integrated analyses of modules with single cell transcriptomes revealed significant overlap between age-associated module genes and both pro-inflammatory and disease-associated microglial clusters. These modules and clusters harbor known neurodegenerative disease genes including *APOE, PLCG2* and *BIN1*. Meta-analyses with published bulk and single cell microglial datasets further supported our findings. Thus, these data represent a well-characterized human microglial transcriptome resource; and highlight age, sex and *APOE*-related microglial immunometabolism perturbations with potential relevance in neurodegeneration.

## Introduction

Microglia are the resident macrophages of the central nervous system (CNS), responsible for clearance of cellular debris and pathological protein aggregates. In the healthy brain they exist in a resting state and can be induced to a reactive state in response to changes in the CNS microenvironment, such as inflammation and neuronal damage^1^. They are fundamental to maintaining brain homeostasis during development, aging and disease, therefore microglial dysfunction could ultimately lead to neurodegeneration^2^. Microglia are integral to the pathophysiology of neurodegenerative diseases, including Alzheimer’s disease (AD) and multiple sclerosis, with chronic inflammation implicated as a contributing factor^3–5^.

Fresh human brain tissue studies are imperative to the characterization of the microglial transcriptome in health and disease; however, accessibility is limited. Although single nuclei studies using frozen tissue provide an easier alternative, recent studies have demonstrated limitations in detecting substantial populations of less abundant cell types^6,7^. Additionally, it was recently reported that many microglial activation genes are expressed in the cytosol and therefore are likely to be missed by single nuclei RNA sequencing (snRNAseq)^8^. Recent single cell studies aiming to characterize microglial gene expression using fresh tissue have highlighted the heterogeneity in microglial phenotypes^9–11^. This has revealed that phenotypic changes are not binary but rather a spectrum of states in which microglia can simultaneously co-exist during transition from resting to more reactive states. Additionally, these different subsets could have specialized functions in brain homeostasis and dysfunction. Thus, it is increasingly important to characterize these heterogeneous subpopulations to understand their roles in health and disease. This could also help facilitate the design of novel therapeutic approaches to target specific subpopulations of cells and modulate their activity^2^.

Utilizing fresh human tissue from neurosurgeries allows us to study the mechanisms of microglial function in living cells. By necessity, this tissue is typically obtained from surgical resection of tumor or epilepsy-affected regions. While there are studies which investigated the effect of glioblastoma (GBM)^12^ or temporal lobe epilepsy^13,14^ on myeloid cells within or surrounding the disease tissue, studies which leverage ‘normal’ tissue from these resections to explore microglial transcriptome are rare^9,11,15^. Despite careful surgical resection to obtain ‘normal’ tissue, there may still be microglial transcriptional changes that occur as a result of nearby disease tissue. Combined analyses of data from multiple available studies can enhance sample size, rigor and reveal common microglial transcriptional signatures robust to differences in tissue source or technique. However, such joint analyses are lacking from the literature. In this study we leveraged the microglial expression data from our neurosurgical samples and that made available by others^9,16–18^ and discovered robust transcriptional changes associated with age, sex and *APOE*.

Microglial expression has been shown to be affected by aging^17,18^, however few studies have investigated the effects of sex and genetic factors on human microglia. Sex differences in microglia have been previously reported in mice, with females being predisposed to harboring more activated microglia than males^19–21^. *APOE*, a lipoprotein of which the ε4 allele (*APOE-*ε4) is a major risk factor for AD and also implicated in other neurodegenerative diseases^22^, is upregulated in disease-associated microglia (DAM) in mice and humans, but downregulated in astrocyte and oligodendrocyte subpopulations^4,6,23,24^. In microglia and neurons, *APOE* interacts with LDL receptors to facilitate endocytosis of cholesterol and phospholipids and modulate lipid homeostasis in the brain^25^. Such studies provide growing support for cell type-specific functions of *APOE*, however, its effects on microglia remain to be fully elucidated. Thereby identifying age, sex and *APOE*-associated pathways in microglia will provide greater insight into the functions of specific microglial subsets in relation to these risk factors. Inter-individual variability and diversity in functional states makes targeting specific microglial subsets in disease challenging for modulating these cells^2^. Identifying the mechanisms regulating microglial homeostasis and activation can allow us to manipulate these cells for therapeutic purposes.

In this study, we leveraged both bulk and single cell approaches to provide a comprehensive characterization of the adult human microglial transcriptome. We obtained fresh intraoperative neurosurgical brain tissue and isolated an enriched population of microglial cells to investigate transcriptional changes associated with age, sex and *APOE*-ε4 in bulk microglia and further explored these in single microglial cells. Our findings support age-, sex- and *APOE*-related microglial transcriptome changes involving lipid and carbohydrate metabolic pathways and implicate microglial immunometabolism perturbations relevant to neurodegenerative diseases.

## Results

To uncover microglial transcriptional profiles and their associations with age, sex and *APOE*, we performed microglial cell-type specific and single cell RNA sequencing (scRNAseq) studies in fresh human brain tissue. We obtained neurosurgical tissue unaffected by the primary disease process from 19 human donors (**Supplementary Figure S1**). Microglia were isolated by CD11b^+^ microbead selection followed by FACS sorting of cells expressing the CD11b^+^/CD45^intermediate^ microglial signature to acquire a more purified population. These samples underwent bulk microglia RNAseq, with subsets of these and additional samples also undergoing 10x scRNAseq (n=5) and bulk tissue RNAseq (n=9) (**Figure 1A**; **Supplementary Table S1**). Validation of sorted microglia using qPCR showed the expected *CD11b*^+^/*CD45*^intermediate^/*P2RY12*^+^ microglial signature^2^ with no expression of other cell type markers, indicating that we isolated a highly enriched microglial population (**Supplementary Figure S2**).

**Figure 1.**
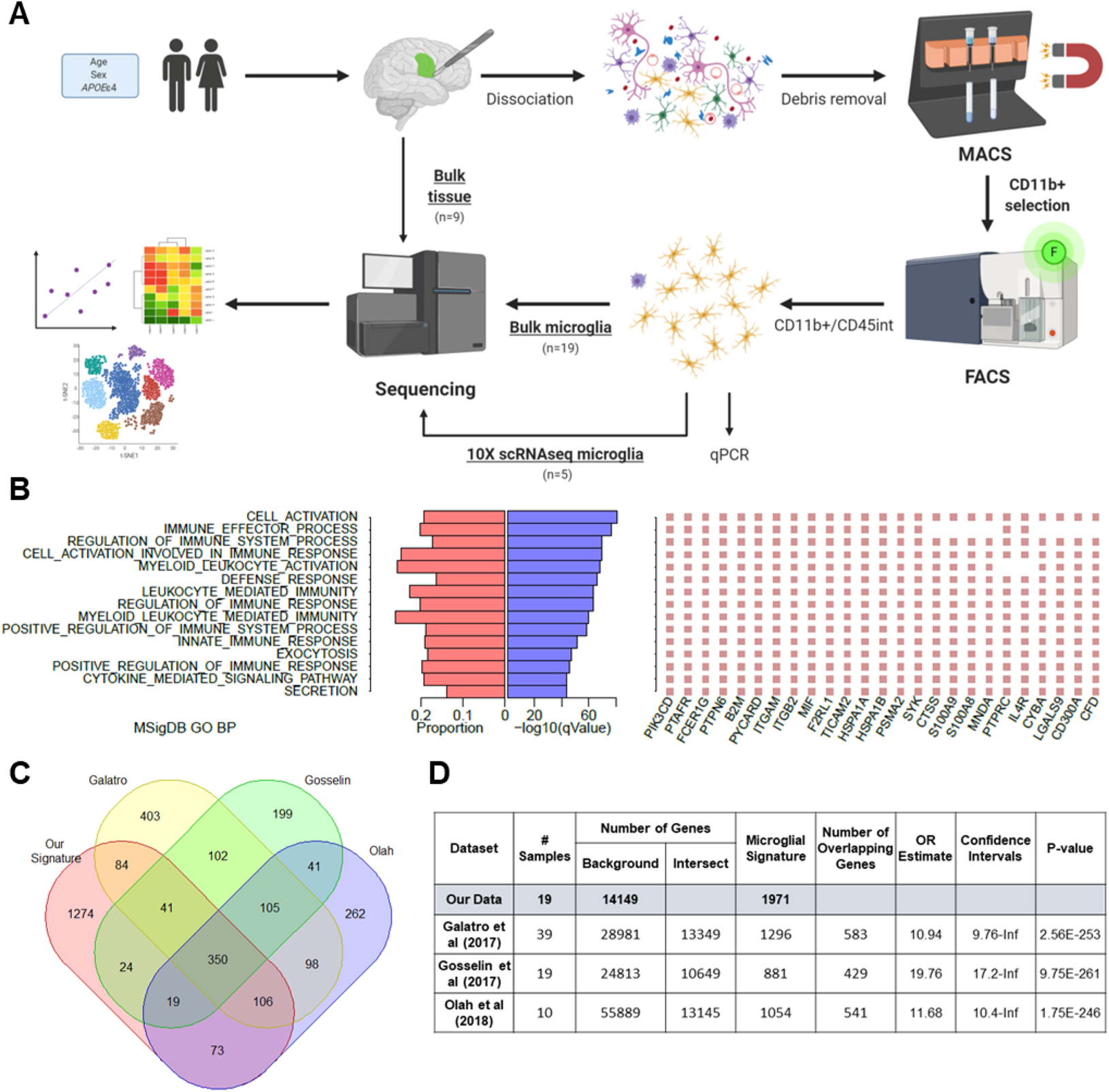
Characterization of our core human microglial signature. (A) Schematic illustrating our experimental approach for isolating microglial populations from fresh brain tissue and data analyses. [Created with BioRender.com] (B) MSigDB GO terms enriched in our microglial signature genes and top 25 genes for each. (C) Venn diagram showing number of overlapping genes between our microglial signature and those previously reported from Galatro et al (2017), Gosselin et al (2017) and Olah et al (2018). (D) Hypergeometric tests of overrepresentation showing overlap with the published signatures.

### Identification of a core human microglial transcriptional signature

To define a core human microglial signature, we calculated log_2_ fold change and q-values of differential expression for each gene between bulk microglia RNAseq data in our study and bulk brain RNAseq data from 7 AMP-AD datasets provided by Mayo Clinic^26^, Mount Sinai Brain Bank^27^ and Rush University Religious Orders Study and Memory and Aging Project (ROS-MAP)^28^ representing 6 brain regions from 515 human samples. Using a cutoff of 4-fold greater expression in our bulk microglia and a q-value threshold of 0.05, we identified 1,971 genes (**Supplementary Table S2**-**4**). These genes were expressed at significantly greater levels in our bulk microglial transcriptome data in comparison to each of the bulk brain transcriptome datasets. Therefore, we considered these 1,971 genes as the core microglial signature in our dataset. This signature comprises several known marker genes, with 12.7% of the genes being BRETIGEA^29^ microglial genes, suggesting that it also likely harbors novel microglial markers of interest (**Supplementary Table S5**). GO enrichment using MSigDB showed that this signature was enriched for genes involved in immune-related and inflammatory response pathways as would be expected, and leukocyte mediated immunity (**Figure 1B**).

To determine the ability of bulk brain tissue data to capture microglial genes, we assessed the expression levels of our microglial signature genes in each of the 7 AMP-AD bulk brain RNAseq datasets. Of the 1,971 microglial signature genes in our study, 37-47% were captured in these bulk brain datasets (**Supplementary Figure S3A-B**). Our microglial signature genes comprised 3.6-4.5% of the expressed bulk brain transcriptome, consistent with prior estimations^6,30^. We next compared bulk microglia RNAseq transcript levels to that obtained from bulk tissue RNAseq of neurosurgical fresh brain tissue samples. Bulk fresh brain tissue does not capture all microglial marker genes, as demonstrated by the low correlation between bulk tissue and bulk microglia data (R=0.46) (**Supplementary Figure S3C, S4, Supplementary Table S6**). This reiterates the need for complementary single cell type data to deconvolute cell type specific expression. We provide the list of microglial signature genes that are also expressed at high levels in bulk brain tissue data (**Supplementary Table S7**), which can serve as a validated resource for microglial signature gene markers in bulk RNAseq datasets.

To determine how the microglial signature in this study compared to previously published signatures, we performed hypergeometric tests of overrepresentation with Galatro, et al. ^17^, Gosselin, et al. ^15^ and Olah, et al. ^18^ studies which had signatures consisting of 881-1296 genes and also with Srinivasan et al.^16^ with a smaller signature of 66 genes. Significant overlap was observed across all datasets, except with Srinivasan^16^ et al., with 350 genes common to ours and the three larger signatures (**Figure 1C-D, Supplementary Table S4, S8**). This comprised several established microglial marker genes, including *P2RY12*, *TMEM119* and *CX3CR1*. The most significant overlap was shared with Gosselin, et al. ^15^ signature [OR=19.6 (17.0-Inf) p=3.8E-261], where 49.7% of their genes were also present in our signature, and 22% of ours in their signature. Gosselin, et al. ^15^ samples were also obtained from neurosurgical tissue resections like our cohort; and are unlike Galatro, et al. ^17^ and Olah, et al. ^18^ samples that were harvested during autopsy. We repeated these analyses focusing on the subset of our samples with matching ages to the comparison groups, where applicable, and still observed significant overlaps in signature (**Supplementary Table S9**), suggesting these findings are robust. Although there appears to be a common set of microglial genes consistent across signatures, each also harbors many unique genes, which could be due to study or individual specific differences.

### Transcriptional profiling of microglia discovers co-expression networks and implicates lipid and carbohydrate metabolism pathways associated with age, sex and *APOE*

We generated gene co-expression networks using WGCNA^31^ to reduce number of tests and increase power to detect genetic associations with age, sex and *APOE*. We identified 7 modules with significant associations (**Figure 2; Supplementary Figure S5; Supplementary Table S10**). Modules ME14 and ME34 associated with age, however, in opposite directions. ME14 was enriched for genes involved in the lipid localization pathway that were upregulated with age (R=0.50, p=0.03) (**Figure 2A-C)**. ME34, enriched for DNA endoreduplication genes, had negative association with both age (R=−0.55, p=0.01) and *APOE*-ε4 (R=−0.50, p=0.03), indicating that microglial transcripts involved in this pathway are downregulated with aging and in *APOE*-ε4 carriers (**Figure 2A**). Several other modules also associated with *APOE*-ε4, in either direction. The only module associated with sex was ME26, which was downregulated in females (R=−0.54, p=0.02), and enriched for genes involved in cholesterol absorption and lipid digestion. This module also had the most significant association with *APOE*, in the positive direction with presence of *APOE*-ε4 (R=0.66, p=0.002) (**Figure 2A,B,E**). Of the *APOE* associated modules, ME23 had the second most significant association (R=−0.61, p=0.006) and was enriched for carbohydrate metabolism genes (**Figure 2A,B,D**). Given recent discoveries in microglial immunometabolism^32–35^, we focused on ME14, ME23 and ME26 that are enriched for lipid and carbohydrate metabolism genes.

**Figure 2.**
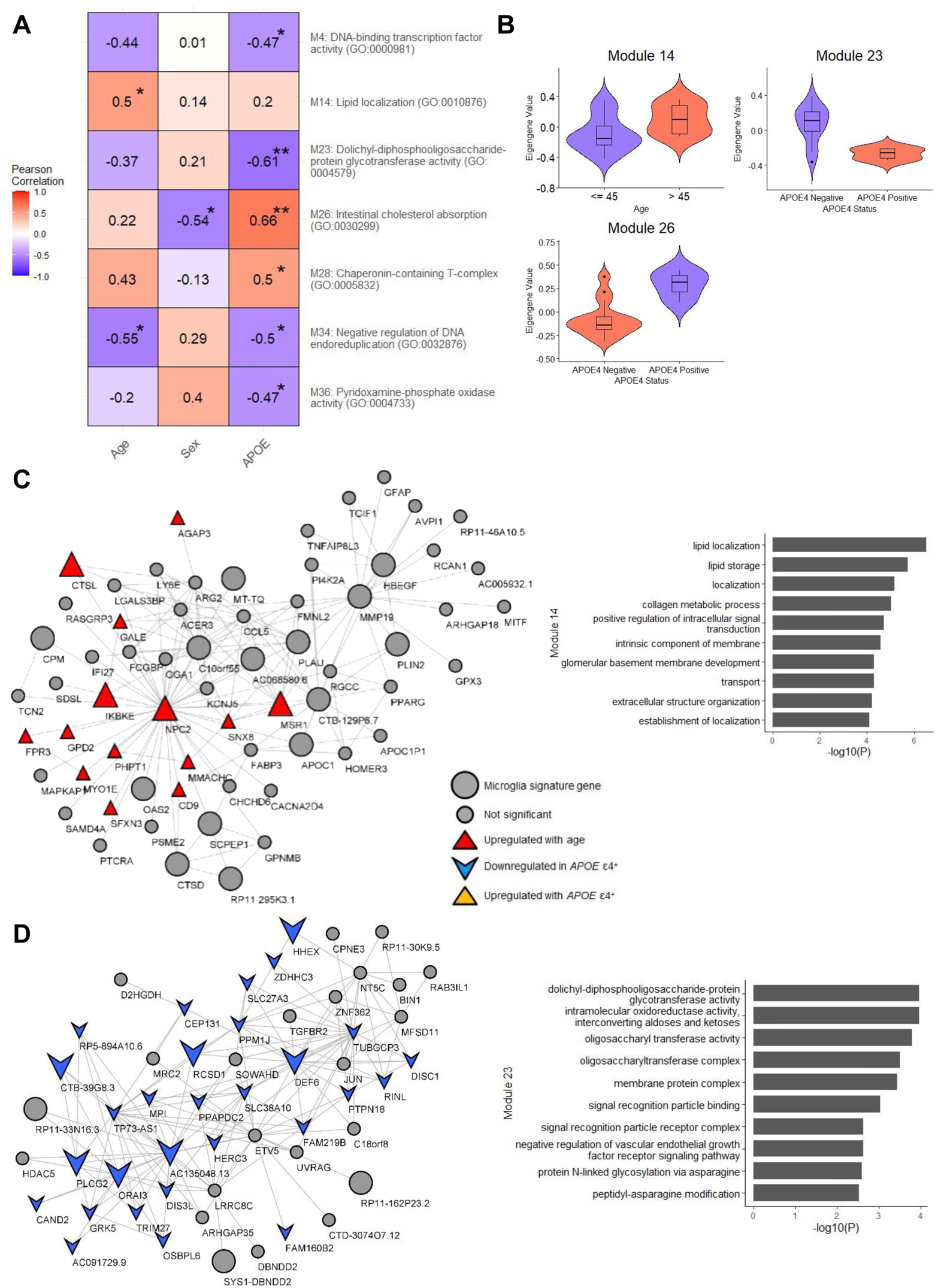
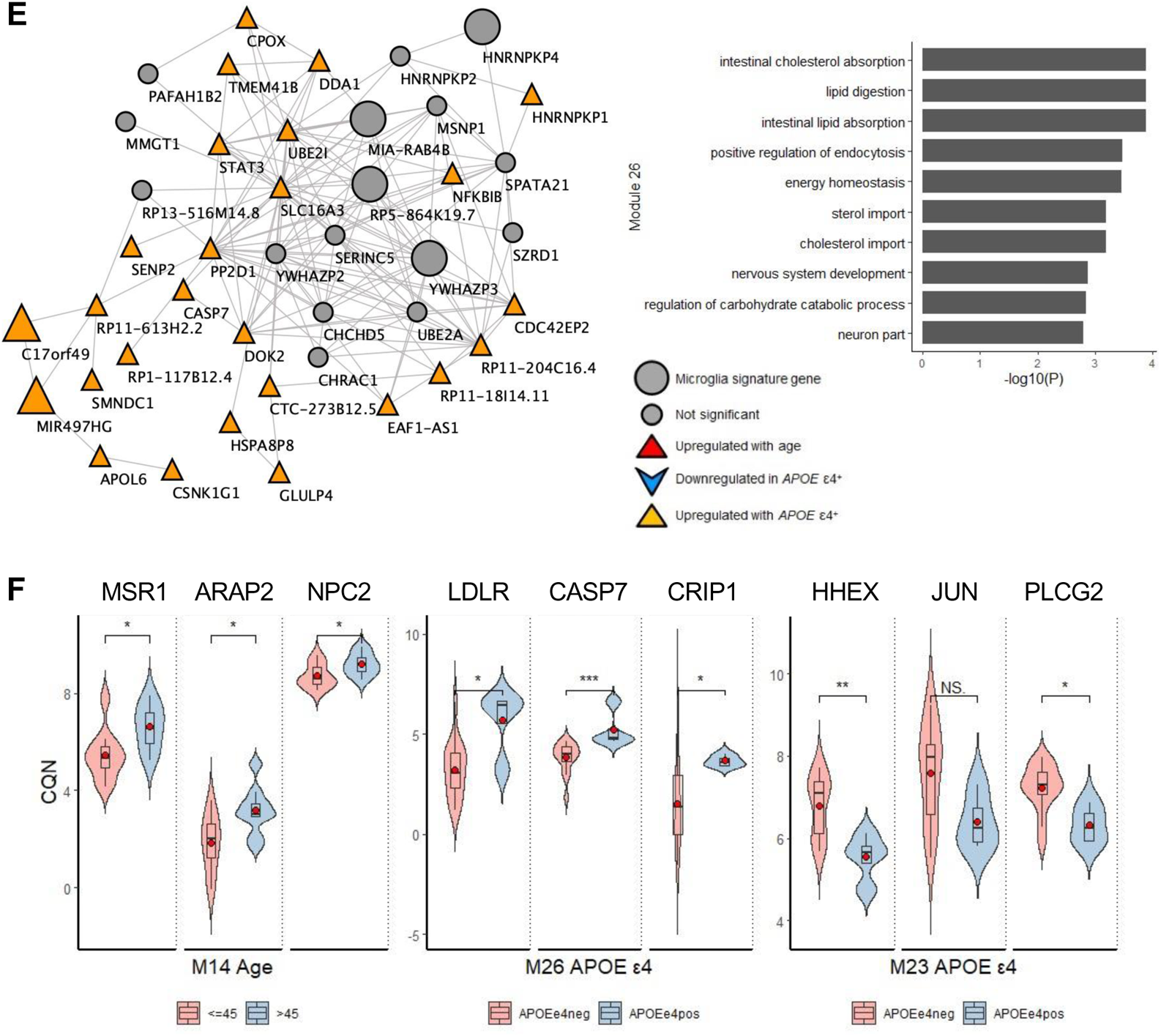
Age, sex and *APOE* ε4 pathway correlations in bulk microglia. (a) Heatmap showing correlation of age, sex and *APOE* ε4 status with WGCNA module eigengenes (MEs) significantly associated (p<0.05) with traits, with top GO terms listed for each module. (b) Module eigengenes stratified by age or *APOE* ε4 status. (c) Module M14 gene co-expression network, with genes of interest highlighted according to the key. Genes upregulated with age shown in red triangle (▴). Bar plot of top 10 significant GO terms (p<0.05) for this module. (d) Module 23 gene co-expression network, with genes downregulated in *APOE* ε4 carriers shown in blue arrow (▾). Bar plot of top 10 significant GO terms (p<0.05) for this module. (e) Module 26 gene co-expression network, with genes upregulated in *APOE* ε4 carriers shown in orange triangle (▴). Bar plot of top 10 significant GO terms (p<0.05) for this module. (f) Violin plots showing expression of key genes in modules, stratified by age or *APOE*. * p < 0.05; ** p < 0.01; *** p < 0.001.

ME14 co-expression network (**Figure 2C**) hub genes *NPC2*, *MSR1* and *PLAU* are also microglial signature genes in our study and known to be involved in microglial functions^36–40,41^. Several disease-associated microglial (DAM) markers are also present in this network, including *CD9*, *ARAP2* and *MYO1E*^4,42,43^ that are increased with aging, implicating activated microglial lipid localization pathways in aging (**Figure 2F**). Several genes in this module were also previously linked to neurodegeneration, including *MYO1E*^44,45^, *CTSL*^46^ and *UNC5B*^47,48^. Due to the nature of the neurosurgical tissue, we also compared expression levels of the key module genes between tumor and epilepsy samples (**Supplementary Figure S6**). Out of 18 comparisons we only observed significant differences due to diagnosis in *NPC2, PLAU, APOC1* and *IKBKE*; therefore, it is important to note that some of these associations may be confounded due to the disease state.

Our microglial signature (**Supplementary Tables S2-S4**) had significant overrepresentation of the age-associated ME14 genes (**Supplementary Table S10**) (OR=1.55 [95% CI=1.23-INF], p=0.001), highlighting age-related increases in microglial signature genes. Galatro, et al. ^17^ and Olah, et al. ^18^ also reported age-related microglial signatures. Comparison of ME14 genes revealed significant overlap with Olah, et al. ^18^ (OR=1.34 [95% CI=1.05-INF] p=0.03), but not with Galatro, et al. ^17^ microglial aging signature genes (OR=1.09 [95% CI=0.81-INF] p=0.33).

ME26 cholesterol metabolism pathway genes exhibited reduced expression in males and were elevated in *APOE*-ε4 carriers (**Figure 2A,B**). This module contains known microglial genes *LDLR*, *CD36* and *CRIP1* (**Figure 2E,F)**. Assessment of individual ME26 network genes revealed *C17orf49*, *RP11-589P10.7* and *MIR497HG* to be the only microglial signature genes in this network to be associated with both sex and *APOE* (**Figure 2E**). Other microglial signature genes in ME26 associated with only sex or only *APOE*, suggesting that these traits may have independent effects on expression of some microglial genes. Several *APOE*-associated genes in ME26 were previously implicated in AD, including *CASP7*^49,50^ and *LDLR*^51,52^ (**Figure 2F**).

Carbohydrate metabolism gene enriched module ME23 is downregulated in *APOE*-ε4 carriers (**Figure 2A,B,D**). AD risk genes *BIN1*^53^ and *PLCG2*^54^ are present in this network, which have both been implicated in microglial dysfunction in neurodegeneration (**Figure 2D**).

### Single cell transcriptome reveals specific subtypes of microglia

To uncover distinct microglial subtypes, a subset of sorted microglial samples from neurosurgical brain tissue underwent single cell expression profiling. We obtained 26,558 cells from 5 unique individuals, including one individual who underwent epilepsy surgery and had samples from two brain regions (**Supplementary Table S1**). Analysis of the scRNAseq data from these samples revealed 13 distinct cell clusters which were annotated using established neuronal and glial marker genes from the literature^4,6, 9–11,43,55,56^ (**Figure 3A, Supplementary Table S11**). Myeloid markers (*AIF1*, *PTPRC*, *C1QA*) were detected in all clusters except cluster 12 which expressed oligodendrocyte markers (*PLP1*, *MBP*, *MOBP*). Cluster 9 expressed macrophage-specific markers (*VCAN*, *FCN1*, *CRIP1*, *S100A8*). These two clusters comprised <3% of all cells, indicating that our sorted samples represent a very pure microglial population. Each myeloid cluster had cellular contributions from all samples, albeit with some variability in their proportions, likely due to intrinsic differences between individuals (**Figure 3B, Supplementary Table S12**). For these samples, the most marked difference was observed for macrophages (cluster 9) and homeostatic microglia (cluster 2), which had greater contributions from the mesiotemporal and anterior temporal regions, respectively. This could be due to the proximity of the mesiotemporal sample to the disease-affected region.

**Figure 3.**
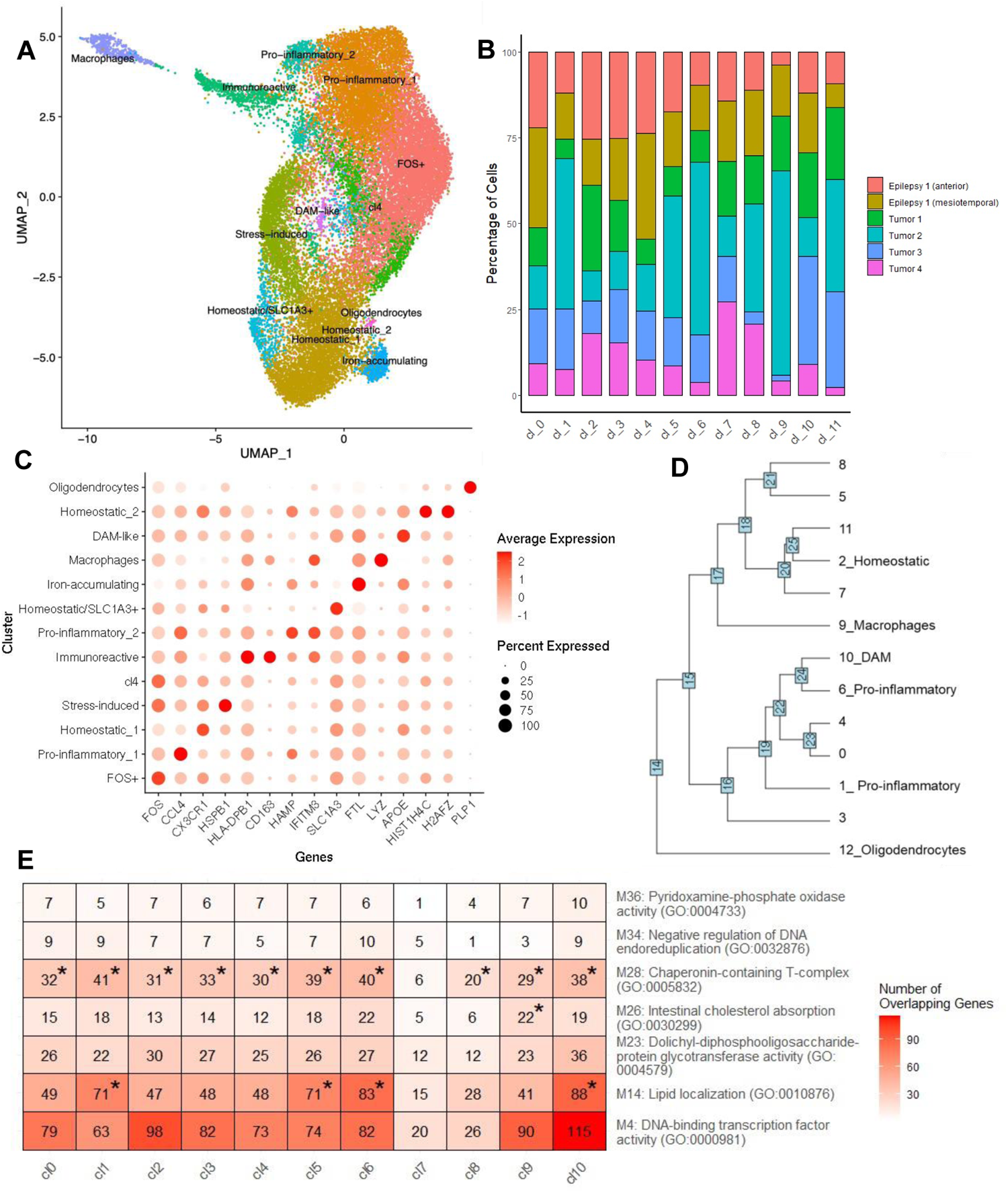
Single cell microglial data. (A) UMAP of clustered cells annotated with putative subtypes using cell type markers from the literature. (B) Stacked bar plot showing the distribution of cells across the clusters. (C) Dot plot showing the expression of key significant module genes across clusters. (D) Hierarchical clustering to highlight relationships between clusters. (E) Hypergeometric distribution of enrichment between module genes and clusters, showing number of overlapping genes. * Represents module genes that were significantly enriched in the cluster (p < 0.05).

We characterized the microglial clusters by their expression of established microglial subtype markers (**Figure 3C, Supplementary Figure S7**) and their most significant marker genes (**Supplementary Figure S8**). Homeostatic (*TMEM119*, *P2RY12*, *CX3CR1*)^10,11,43,56^, pro-inflammatory (*CCL2*, *CCL4*, *EGR2*)^10,11^ and DAM markers (*APOE*, *C1QA*, *C1QB*)^4,9,11,23^ were observed in clusters 2, 1/6 and 10, respectively. Cluster marker genes are defined as those expressed in at least 70% of the cells in the cluster with log fold change > 0.6 and q < 0.05 in comparison to all other clusters. Expression levels of the top marker genes per cluster are shown (**Figure 3C; Supplementary Figure S8; Supplementary Table S13**). Most of these markers are distinct to a single cluster, although some clusters appeared to have similarities in their marker expressions. To define the proximity of their transcriptional profiles, we performed hierarchical clustering of the microglial clusters (**Figure 3D**). We determined that the homeostatic microglia cluster 2 was transcriptionally closest to clusters 7 and 11, which may represent subtypes of homeostatic microglia. Clusters 1 and 6 both expressed chemokines *CCL2* and *CCL4* representative of pro-inflammatory microglia, however cluster 6 was more closely related to DAM, whereas cluster 1 represented a more distinct microglial signature. Cluster 6 highly expressed interferon-related marker *IFITM3* and *ISG15*, also observed in a cluster by Olah et al (2020)^9^, which they defined as an interferon response-enriched subset. These findings highlight different transcriptional profiles for the two pro-inflammatory microglial clusters that may represent distinct activated microglia subtypes. Cluster 3 highly expressed heat shock protein *HSPA1A*, an immediate early gene^57^ reportedly involved in antigen processing^58^, response to stress and injury and exhibiting decreased gene expression in multiple sclerosis patients^59,60^. Several were upregulated in this cluster, suggesting that this may represent cells that underwent dissociation-induced stress^11^. Several of the clusters did not express well known existing cell type markers. Clusters 5/8 and FOS^+^ 0/4 were transcriptionally closest to one another (**Figure 3D**). Cluster 5 has distinct expression of immunoreactive marker *CD163*, which was not observed in other subsets except macrophages. Several *HLA* genes are also highly expressed in this cluster, suggesting that these may be immunoreactive microglia^61^. Cluster 8 marker FTL has recently been used to characterize iron-accumulating microglia^62^. Our findings highlight transcriptional profiles for known microglial clusters, describe the transcriptional proximity of these clusters and suggest that less well-defined clusters could potentially represent novel or intermediate transcriptional states of microglia. Our microglial signature was significantly enriched in more clusters expressing more activated markers (**Supplementary Table S14**), implicating this as the dominant expression profile within our samples. However, there is also enrichment of homeostatic cluster 2, demonstrating that we have not only captured activated cells as might be expected due to the nature of the tissue, but also homeostatic microglia subtypes.

To determine whether the bulk microglial co-expression networks (**Figure 2A,C-E, Supplementary Figure S5**) were representative of microglial subtypes, we performed enrichment analyses of the module genes within the myeloid clusters with sufficient cell numbers (**Figure 3E; full enrichment statistics provided in Supplementary Table S15**). Age-associated co-expression network ME14, implicated in lipid metabolism, was significantly enriched in pro-inflammatory (cluster 6) and DAM (cluster 10) clusters. Genes within module 28, which was significantly upregulated with *APOE-*ε4, had statistically significant enrichment in all clusters except cluster 7. There was no statistically significant enrichment for any of the other microglial modules that had significant age, sex or *APOE* associations, suggesting that these factors may have ubiquitous effects on most microglial subtypes. Some of the remaining microglial co-expression networks had distinct patterns of cluster enrichment (**Supplementary Figure S9**), suggesting that some but not all networks could be representative of distinct microglial subtypes.

### Meta-analyses with published datasets support age, sex and *APOE* associations

We identified three bulk microglial^16–18^ and one single cell microglial^9^ transcriptome study with available data that could be analyzed with our own. Each bulk microglia transcriptome dataset was individually processed through the same MAPR-seq pipeline^63^ for quality control to minimize variability due to data processing. Meta-analysis of WGCNA results was performed by coercing the external dataset co-expression networks onto our existing co-expression networks. The forest plot in **Figure 4A** highlights the individual and combined associations for networks of interest across the datasets where age, sex or *APOE* information was available (**Supplementary Table S16**). For module ME14 associations with age, Galatro et al (2017) exhibited a similar direction of effect (R=0.30, P=0.06), whereas Srinivasan et al (2020) did not show any association (R=−0.003, P=0.99), likely due to a higher median age of individuals in that study. Meta-analyses of these two datasets with ours showed association of ME14 with age (R=0.25, P=0.05). ME26 showed lower levels in males in all but Olah et al^18^. When meta-analyzed, sex-associated ME26 genes were significantly down-regulated in males (R=−0.30, P=0.007), supporting our finding. This module was also significantly associated with *APOE* e4 carriers in the Srinivasan dataset (R=0.42, P=0.04), as in ours (R=0.57, P=0.01) with significant meta-analysis association (R=0.49, P=0.001). Likewise, Srinivasan data was inversely correlated with *APOE* for ME23 (R=−0.38, P=0.07), as is our data (R=−0.58, P=0.009), with significant meta-analysis results (R=−0.0.47, P=0.0.002). The consistent direction of association between most datasets and ours is also evident when hub genes from the modules are plotted by the analyzed variable (**Supplementary Figure S10**). These consistent results across multiple datasets provide support for our findings.

**Figure 4.**
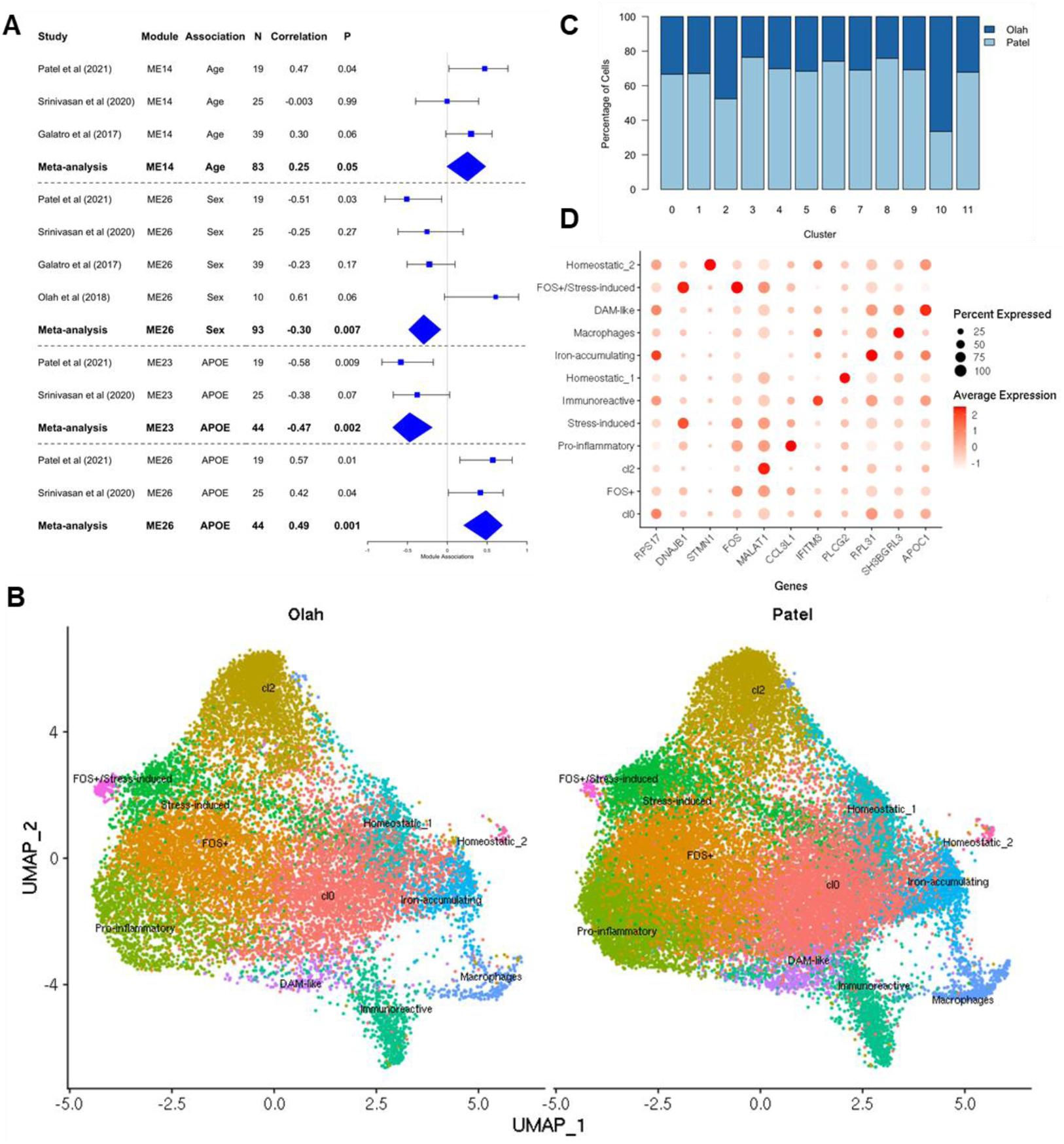
Meta-analysis with published datasets. (A) Forest plots of module eigengene correlations across individual bulk microglial datasets. Results from meta-analyses of relevant samples are shown in bold. (B) UMAPs of Olah et al’s (2020) (left) and our (right) single cell data, split by dataset. Samples were integrated prior to clustering and downstream analyses. Cells from each dataset contribute to all clusters. (C) Stacked bar plot showing the distribution of cells across each of the clusters. (D) Dot plot showing the expression of top cluster marker genes across all clusters.

We subsequently integrated microglial single cell data from Olah et al (2020)^6^ (n=9, 15,819 cells) with our samples (n=6, 26,558 cells) to compare these single cell datasets. There were 11 integrated-clusters when both datasets were analyzed together. The UMAP in **Figure 4B** is split by study and shows relatively even contributions to each cluster from both datasets, also observed in the stacked bar plot in **Figure 4C**, demonstrating slightly greater proportions from our dataset commensurate with our greater cell numbers. One exception is the integrated-cluster 10 that has a greater number of cells from Olah et al. This cluster has similar expression patterns to our original FOS^+^ cluster 0. Hypergeometric tests of enrichment showed significant overlap between many of the original and integrated clusters, showcasing the high congruence between our data and Olah et al.^9^ (**Supplementary Table S17**). To further identify where our individual cells clustered when combined with the Olah dataset, we overlaid the cell IDs from the original clusters to the integrated dataset and calculated the overlap (**Supplementary Table S18**). Greatest cellular overlaps were observed between the original clusters and integrated-clusters with the same subtype features. Many integrated-clusters had the identities observed in the original cluster, as evidenced by the dot plot (**Figure 4D**) which shows some of the same top marker genes depicted in **Figure 3C**. We again observed enrichment of our microglial signature in several integrated single cell clusters, many of which highly expressed markers of activated microglia (**Supplementary Table S19**). Finally, we compared the microglial subtype marker genes from a published single cell^11^ and single nucleus^6^ study for many of the clusters from our single cell data (**Supplementary Table S20**), with greater enrichment for like-subtypes from the former dataset and ours. Collectively, these findings demonstrate the robustness of the microglial subtypes we identified and provide detailed characterization of these in both our and integrated single cell datasets.

## Discussion

Given their critical functions in maintaining homeostasis in the central nervous system (CNS) in health and their multifaceted roles during neurological diseases^2,3^, understanding the biology of microglia and characterizing microglial subtypes is essential. Large scale studies in bulk brain tissue^26–28^ have been instrumental in establishing transcriptional profiles in health and neurodegenerative diseases. Although these studies yielded information on brain expression signatures and uncovered perturbed pathways and molecules implicated in Alzheimer’s disease and other neurological disorders^64–67^, they are limited in their ability to provide cell-type specific transcriptional outcomes, especially for less abundant CNS cells such as microglia^30^. Analytic deconvolution approaches began to leverage these bulk tissue transcriptome datasets to estimate cell-type specific expression profiles^29,30^, but the accuracy of these methods relies on the availability of high-quality single cell-type datasets. Such microglia-specific transcriptome datasets are gradually emerging^9,15,17,18^, although the numbers of unique samples assessed remain limited given the arduous nature of collecting fresh human brain tissue. Additionally, comparative assessment of bulk brain vs. single cell-type bulk microglia vs. single-cell microglia studies are still rare^9,16,68^. To our knowledge there are no studies that evaluate human microglial transcriptome using all three approaches, as in our study. Further, investigations on effects of genetic and other factors on microglial transcriptional signatures in humans is likewise sparse, with the exception of age-related effects assessed in a few studies^15,17,18^. Finally, unlike in bulk tissue studies^29,64–67^, microglia-specific co-expression networks, their molecular signatures and functional implications have not been evaluated.

In this study, we sought to overcome these knowledge gaps by characterizing the transcriptome of sorted bulk and single-cell microglial populations isolated from fresh human brain tissue. We identified a robust microglial signature comprising 1,971 genes enriched for immune-related functions. These signature genes were selected due to their consistently higher expression levels in our sorted bulk microglial transcriptome in comparison to 7 different bulk brain tissue datasets from 6 different regions^26–28^. We also compared sorted bulk microglia to bulk fresh brain tissue and identified transcripts that are expressed in both. The microglial signature genes that are also reliably detected in bulk brain tissue represent a validated list of microglial markers that can be utilized in bulk brain tissue transcriptome analytic deconvolution studies^29,30^.

Our microglial signature significantly overlapped with other signatures from bulk microglia previously reported by Galatro, et al. ^17^, Gosselin, et al. ^15^ and Olah et al.^18^, implicating a core set of genes consistently expressed in this cell type. However, there were additional genes unique to each signature, likely to be driven by factors such as patient demographics or study differences. Galatro, et al. ^17^ and Olah et al.^18^ both also reported age-related microglial expression signatures. We found significant overlap of our age-associated microglial gene expression module ME14 genes with the latter, which was also enriched for our microglial signature. This indicates that bulk microglial profiles can effectively capture genes affected by aging in microglia.

We leveraged the co-expression network structure of sorted bulk microglia to further explore whether microglial subsets were associated with age, sex or *APOE*-ε4. To our knowledge sex-differences in microglial transcriptome were previously studied only in mice^19–21^, however *APOE* genotype-specific microglial interactions with amyloid plaques have been previously observed in mice^20,69^ and humans^70^. We identified two network modules associated with age, one with sex and six with *APOE*-ε4. We observed that two modules, ME14 that is positively associated with increased age; and ME26 that is positively associated with both *APOE*-ε4 and female sex, were both enriched for lipid metabolism biological terms^32–34^. Module ME14 included genes involved in lipid localization and storage pathways (*PLIN2, IL6, LPL, MSR1, ENPP1, PPARG, PTPN2, SOAT1, IKBKE*) and ME26 had lipid digestion/cholesterol transport pathway genes (*CD36, LDLR*). Both modules harbored known microglial genes (*LDLR*, *CD36, CRIP1, NPC2*, *MSR1, PLAU)* and those that are included in our microglial signature (*PLIN2, IL6, MSR1, SOAT1, IKBKE, NPC2, PLAU*).

Comparing the sorted bulk microglial network modules to scRNAseq microglial clusters, we determined that ME14 genes were significantly over-represented in pro-inflammatory cluster 6 and disease-associated microglia (DAM) cluster 10. In our study, DAM cluster 10 included *APOE, APOC1, ASAH1* and *CTSD*. Of these *APOE*^22,33,71^*, APOC1* and *ASAH1*^72^ are involved in lipid metabolism and neurodegenerative diseases. *APOE*^4,5,23^*, APOC1*^23^ and *CTSD*^4^ were also signature genes in mouse models of neurodegenerative diseases^4,5^ or aging^23^. Our pro-inflammatory cluster 6 also included genes associated with mice microglial neurodegenerative (*FTH1*^4^) or aging signatures (*CCL4*^23^), as well as *IFITM3*^34^*, GOLGA4*^34^, previously shown to be upregulated in aging lipid droplet accumulating microglia^34^. Our findings that integrate human sorted bulk RNAseq and scRNAseq data, support a model where aging human microglia transition to a pro-inflammatory and disease-associated transcriptional profile which is also associated with perturbations in lipid metabolism in these cells.

There is increasing evidence that tightly controlled lipid metabolism is essential to the functions of microglia during development and homeostatic functions of adulthood and may be disrupted in aging and disease^32,33^. The complex interactions between microglial lipid metabolism and its cellular functions rely on lipid sensing by microglial receptors such as CD36 and TREM2 and uptake of lipids, including LDL and APOE^32,33^. These interactions are necessary for microglia to become activated and perform functions including phagocytosis of myelin^73^ and misfolded proteins like amyloid ß^74^, cytokine release, migration and proliferation^32,35^. Studies primarily focused on *in vitro* and animal models suggest disruption of the microglial immunometabolism and assumption of a pro-inflammatory phenotype with aging^23,34,75,76^ and diseases including multiple sclerosis (MS) and Alzheimer’s disease^4,5,77^. Interestingly, microglial lipid droplet accumulation has been demonstrated under all these conditions^32–34,73^ and lipid droplet accumulating microglia in aging mice were shown to have a unique transcriptional state^34^. Our findings in sorted cells from fresh human brain tissue provide transcriptional evidence for immunometabolism changes and pro-inflammatory phenotype with microglial aging, thereby contributing essential complementary data from humans for this cell type.

Besides module ME14, we determined that ME26 is also enriched for lipid metabolism genes. ME26 module expression is higher in both *APOE*-ε4 and female sex, however we note that in our sorted bulk microglia RNAseq samples, there were no male *APOE*-ε4 carriers. Therefore, the distinct influence of sex and *APOE* on the expression of this module remains to be established. *APOE*-ε4, a major risk factor for Alzheimer’s disease, has the lowest lipid binding efficiency compared with other *APOE* isoforms^32^. Increased cholesterol accumulation has been reported in both iPSC-driven astrocytes from *APOE*-ε4 carriers^78^ and in Apoe-deficient microglia^73^. These findings collectively support a role for *APOE*-ε4 associated microglial transcriptional changes and disrupted cholesterol metabolism. Using our sorted microglia RNAseq data, we identified five additional modules that associate with *APOE*-ε4, one in a positive direction (ME28) and four negatively (ME4, ME23, ME34, ME36). Of these, module ME23 had the second most significant *APOE*-ε4 association after ME26. Interestingly, ME23 was enriched for carbohydrate metabolism biological processes, which are also tightly regulated in microglia^35^. Module ME23 harbors known AD risk genes *BIN1* and *PLCG2*, where the latter is a microglial gene that modulates signaling through *TREM2*^79^ and also a hub gene in this module. ME23 genes *BIN1, JUN* and *TGFBR2* were found to be reduced in a mouse microglial neurodegenerative phenotype gene signature^5^. These findings further demonstrate the consistency of our human microglial data with that from mouse models and supports perturbed microglial immunometabolism as a potential pathogenic mechanism in neurodegeneration. Importantly, using available transcriptome data from three other bulk microglia datasets of Galatro et al (2017)^17^, Olah et al (2018)^18^ and Srinivasan et al (2020)^16^, we performed meta-analyses with our data thereby providing results from a collective sample size of 93, 83 and 44 fresh tissue samples for respectively, sex, age and *APOE* associations. We observed that the associations in our data replicated in most of the other available datasets and the meta-analyses improved the significance of most associations. These results represent the largest collective bulk microglial associations from human fresh brain tissue to our knowledge and provide further support for our findings.

In addition to analyzing gene expression modules from sorted bulk microglia, we also identified microglial clusters from sorted microglial scRNAseq data. To our knowledge, there are only two prior publications of scRNAseq characterizations on human microglia^9,10^. Masuda et al.^10^ analyzed 1,602 microglia isolated from 5 control and 5 MS patient brains, compared their findings to those from mice demonstrating clusters that are common and others that are species-specific. Olah et al. assessed 16,242 microglia from 17 individuals and characterized subclusters of microglia from patients with mild cognitive impairment, AD and epilepsy^9^. Our scRNAseq dataset is from 5 unique individuals comprising 26,558 cells, 99.98% of which have myeloid markers. We identified microglial clusters that share characteristics of those previously reported in mice^4^ and humans^9,70^, such as DAM. We also uncovered clusters that had not been previously characterized in the literature, including cluster 7, exhibiting high expression levels of astrocytic marker gene *SLC1A3*. Microglial expression of *SLC1A3* was previously shown to occur in mice and humans especially in disease states^80–82^. We also leveraged these scRNAseq data to further characterize the sorted bulk microglial expression modules. Integration of our samples with Olah et al.^9^ allowed us to explore microglial subtypes at single cell resolution with increased power. We found that even in this larger dataset, our microglial signature was similarly enriched in the more activated subtype clusters, demonstrating the robustness of this signature across datasets. The homogeneity in clustering highlighted shared expression patterns between cells from both datasets, providing further support for the putative microglial subtypes we identified in our current study. Hence our microglial scRNAseq data contribute further to the emerging single cell landscape of this cell type.

Activated microglia are a hallmark in both brain cancers and temporal lobe epilepsy in response to the inflammation^12–14,83,84^. We acknowledge that the transcriptome data generated in this study is sourced from grossly normal tissue obtained during resection of brain tumor or epilepsy regions. Despite careful resection to exclude disease tissue, it is possible that the grossly normal tissue transcriptome may still be influenced by nearby pathology. Therefore, we cannot rule out the possibility that the activated microglial subtypes we identify may be due to the nature of the tissue. Indeed, sex-specific expression differences have been observed in glioma-activated microglia^84^. Darmanis et al^12^ investigated the effect of glioblastoma tumors on CNS cell types and surrounding tissue, and found that peri-tumor myeloid populations were primarily pro-inflammatory microglia compared to macrophages within the tumor core^12^. In temporal lobe epilepsy, two distinct microglial phenotypes have been identified with microglia present in sclerotic areas with few neurons expressing markers of activation, including anti-inflammatory cytokine IL10^13,14^. The other phenotype occurs transiently following a seizure, with secretion of interleukins CXCL8 and IL1B mediated by the NLRP3 inflammasome^13^. Nonetheless, we did not observe significant differences in most genes we highlighted here based on tumor vs. epilepsy diagnosis (**Supplementary Figure S6**). There is also a large cluster of homeostatic microglia detected in our samples, indicating that not all cells are activated. Further, the consistency of our findings with other single cell^9^ or bulk microglia^16–18^ data supports the notion that the microglial signatures and associations we identified in our study are applicable to microglia biology in general rather than being disease-specific.

We recognize that our study has several limitations, primarily owing to the difficulty in obtaining high quality neurosurgical brain tissue, which leads to limited sample size and variability in tissue, diagnoses and patient demographics. Even though we have utilized grossly normal tissue surgically separated from disease tissue, the samples are from epilepsy and various brain tumor patients representing multiple diagnoses. We isolated microglia using an approach which should minimize activation however we cannot definitively rule out stress-induced transcriptomic changes during isolation. Despite these caveats, we could identify microglial co-expression modules and subclusters with multiple features that are consistent with prior publications from model systems^4,5,23,34^. Our scRNAseq clusters have contributions from both tumor and epilepsy samples, suggesting that our findings are unlikely to be driven by any one diagnoses. Furthermore, there are few published studies using fresh brain tissue to study microglia and thus our integrated single cell data showing a homogenous cluster of microglia highlights the robustness of the methodology. Finally, our joint and meta-analysis using all available datasets enhance the rigor of and support our conclusions.

In summary, our study on sorted bulk microglia RNAseq and scRNAseq from fresh brain tissue yield several key findings. We identify a microglial gene signature from sorted bulk microglia, characterize its expression in bulk brain RNAseq across 7 datasets comprising 6 regions, in bulk fresh brain RNAseq and in microglial scRNAseq subtype clusters. This signature provides a well-characterized resource which can be utilized in analytic deconvolution studies of bulk transcriptome data^29,30^. We uncovered microglial gene expression modules associated with age, sex and/or *APOE*-ε4. Modules with age and *APOE*-ε4 associated transcriptional changes implicate microglial lipid and carbohydrate metabolism perturbations and microglial activation. Microglial scRNAseq data highlight the transcriptional complexity of this cell type, reveal both known and novel cell types and demonstrate utility of this data in characterizing sorted bulk RNAseq data. These findings provide support for the emerging microglial immunometabolism^32,35^ pathway as a plausible therapeutic target in aging-related disorders; and provide a well-characterized human transcriptome resource for the research community on this cell type with central roles in health and disease^1^.

## Methods

### Patient Samples

Fresh human brain tissue was obtained from patients undergoing neurosurgical procedures for epilepsy or tumor resection. Tissues determined to be grossly unaffected by the primary disease process were utilized for the present study (**Supplementary Figure 1**). Patient samples were transported from the operating room to the laboratory in 1X DPBS (Thermofisher; 14287080) for processing within 1-2 hours of resection. Human tissue was collected with informed consent prior to surgery and all procedures were approved by the Mayo Clinic Institutional Review Board and are HIPAA compliant.

### Tissue Dissociation

Tissue was dissected to remove necrotic tissue, white matter and excess vascular tissue, to retain only cortical grey matter. The remaining tissue was cut into sagittal slices and weighed before being processed using the Adult Brain Dissociation Kit (Miltenyi; 130-107-677) as per the manufacturer’s protocol. Debris removal (Miltenyi; 130-109-398) and red blood cell lysis (Miltenyi; 130-094-183) were also performed. All procedures were carried out on ice. The resulting homogenate was filtered through a 70µm filter before proceeding.

### Magnetic-Activated Cell Sorting (MACS)

The cell suspension was first enriched for CD11b^+^ cells by incubating with anti-CD11b microbeads (Miltenyi; 130-049-601 clone M1/70) for 15 minutes according to manufacturer’s recommendation. This was then washed with PB buffer (0.5% BSA, 1X PBS Ca^2+^/Mg^2+^ free pH 7.4) and filtered through a 70µm cell strainer before being applied to a large separation column (Miltenyi; 130-042-401) in a QuadroMACS separator magnet (Miltenyi; 130-090-976). The CD11b^+^ fraction was collected and resuspended in sterile filtered FACS buffer (1X PBS Ca^2+^/Mg^2+^ free, 0.5% BSA, 2% FBS, 3mM EDTA) for antibody staining.

### Fluorescence-Activated Cell Sorting (FACS)

MACS sorted CD11b^+^ cells subsequently underwent FACS sorting to further purify the microglial population. The cell suspension was incubated in Human TruStain FcX blocking solution (1:20, Biolegend; 422302) at room temperature for 10 minutes. Subsequently, cells were stained with anti-CD11b PE/Cy7 (1:100, Biolegend; 101206, M1/70) and anti-CD45 Alexa Fluor 647 (1:100, Biolegend; 304056, HI30) antibodies for 30 minutes on ice. Following two washes with FACS buffer, SYTOX Green viability dye (1:1000, ThermoFisher; S7020) was added for an additional 20 minutes. Single cell suspensions were filtered through a 40µm cell strainer (Falcon; 352235) before sorting on a BD FACS Aria II (BD Biosciences). CD11b^+^/CD45^intermediate^/SYTOX green^-^ cells were sorted directly into FACS buffer. Independently CD11b and CD45 are not microglial specific markers, however gating cells with a CD11b+/CD45^intermediate^ signature allowed us to differentiate microglia from macrophages which are expected to be CD45^high^. An example of our FACS gating strategy is provided in **Supplementary Figure S2A**.

### RNA Isolation and Sequencing

RNA from sorted microglial cells was isolated using the miRNeasy Serum/Plasma Kit (QIAGEN; 217184) and quantified on the Agilent BioAnalyzer 2100. cDNA libraries were generated using SMARTSeq2 v4 and Nextera Low Input Library Prep Kit. Samples were multiplexed and sequenced on the Illumina HiSeq 4000.

RNA from frozen bulk tissue was isolated using Trizol and chloroform, followed by DNase and clean up using the RNeasy Kit (QIAGEN; 74106). Libraries were generated using the TruSeq Stranded mRNA Library Prep Kit. Samples were multiplexed and sequenced on the Illumina HiSeq 4000. Base-calling of all sequence data was performed using Illumina’s RTA v2.7.7.

### 10X Single Cell 3’ v3 Library Preparation of Sorted Microglia

Viability of MACS plus FACS sorted cells was assessed by Trypan blue (Gibco; 15250061) exclusion and cell density was determined using a hemocytometer prior to adjustment to target 4000-5000 cells. Cells were loaded onto a 10X Chromium chip and run on the GemCode Single Cell Instrument (10X Genomics) to generate single cell gel beads-in-emulsion (GEMs). Single cell RNA-seq libraries were prepared using the Chromium Single Cell 3’ Gel Bead and Library Kit v2 and v3 (10X Genomics; 120237) and the Chromium i7 Multiplex Kit (10X Genomics; 120262) according to the manufacturer’s instructions. Quality of cDNA libraries was determined using a BioAnalyzer 2100 DNA High Sensitivity assay (Agilent; 5067-4626) prior to sequencing one per lane on an Illumina HiSeq 4000.

### Validation with Quantitative Real-Time PCR

Total RNA was extracted from sorted cells using the miRNeasy Serum/Plasma Kit (QIAGEN; 217184). Concentration and quality were assessed using the Agilent BioAnalyzer RNA 6000 Pico Kit (Agilent; 5067-1514). RNA was normalized to 0.5ng/µl for cDNA synthesis using the SuperScript IV VILO Master Mix (ThermoFisher; 11756050). TaqMan PreAmp Master Mix (ThermoFisher; 4391128) was used to pre-amplify the cDNA, followed by TaqMan Universal PCR Master Mix (ThermoFisher; 4304437) with the following gene expression probes: *MOG, AQP4, THY1, PTPRC, ITGAM, P2RY12, PECAM1, CD34, GAPDH* (ThermoFisher; Hs01555268_m1, Hs00242342_m1, Hs00174816_m1, Hs04189704_m1, Hs00355885_m1, Hs00224470_m1, Hs01065279_m1, Hs02576480_m1, Hs99999905_m1). RT-qPCR was performed on a QuantStudio 7 Flex Real-Time PCR System (ThermoFisher) using a relative standard curve to quantify gene expression.

### Validation with Immunocytochemistry

Cultured cells were fixed with 4% paraformaldehyde (PFA) overnight at 4°C and blocked with blocking solution (10% BSA, 5% normal goat serum and 0.1% Triton-X). Fixed cells were stained with anti-TMEM119 (1:100, Biolegend; 853302) extracellular primary antibody with Goat anti-mouse IgG secondary antibody conjugated to Alexa-488 (1:100, Abcam; ab150113). Nuclei were stained with 1µg/ml DAPI (1:1000, ThermoFisher; 62248) before mounting with AquaPoly Mount (Poly Sciences, 18606-20). Images were acquired with a Zeiss LSM880 Confocal microscope using a Plan-Apochromat 20x magnification and 0.8 objective at 1024 by 1024 pixels with a 0.5 microsecond pixel dwell time.

## Data Analysis

### Bulk Microglia RNA-seq Processing

The MAPR-Seq pipeline^63^ was used to align reads to human reference genome hg38 using STAR^85^ and count reads using featureCounts^86^. FastQC was used for quality control (QC) of raw sequence reads, and RSeQC was used for QC of mapped reads. Quality measures were examined including base calling quality, GC content, mapping statistics and sex check to ensure consistency between the recorded and inferred sex from expression of chromosome Y genes. Raw read counts were normalized using Conditional Quantile Normalization (CQN) to generate log_2_ scaled expression values via the Bioconductor package cqn, accounting for sequencing depth, gene length and GC content. Normalized CQN expression values were assessed using Principal components analysis (PCA) to identify and remove outliers, defined as greater than 4 standard deviations from the mean of the first two principal components. In addition, RPKM (reads per kilo bases per million) values were calculated.

### Identification of a Core Microglial Signature from Bulk Microglia Data

To define a core microglial signature, we compared our bulk microglia data to cognitively normal control samples from the AMP-AD bulk tissue transcriptome data from 7 different datasets representing 6 brain regions (Synapse ID: syn2580853); Mayo Clinic^26^ (cerebellum and superior temporal gyrus), Mount Sinai Brain Bank^27^ BM10 (frontal pole), BM22 (superior temporal gyrus), BM36 (parahippocampal gyrus), BM44 (inferior frontal gyrus) and Rush University Religious Order Study-Memory and Aging Project (ROS-MAP)^28^ (dorsolateral prefrontal cortex). Raw gene counts and metadata (see Acknowledgements) were obtained from the AMP-AD RNAseq Harmonization study which had performed alignment and processing of all datasets and brain regions through a consensus pipeline^87^. Samples were removed that had inconsistent sex between that indicated in metadata and that inferred from RNAseq expression; a RIN < 5; were identified as gene expression outliers based on principal component analysis (PCA) (> 4 standard deviation (SD) from mean PC1 or PC2), or missing metadata. In addition, duplicates (lowest read count sample removed) and those with rRNA (>5%) were removed from the MSBB datasets. Furthermore, samples not meeting neuropathological criteria as Alzheimer’s disease (AD)^88^ or control were excluded. To generate the microglial expression signature, only control samples from the AMP-AD datasets were included. Raw read counts were normalized using Conditional Quantile Normalization (CQN). Log_2_ fold change and q-values between each bulk tissue brain region and the bulk microglia profiles were calculated for each gene via linear regression using log_2_(RPKM) without correction for covariates. Genes were filtered using a cutoff of 4-fold greater expression in bulk microglia compared to each bulk tissue region and q < 0.05. Genes that passed these criteria and were significant in comparisons with all 7 bulk brain datasets determined the microglial signature. These signature genes were assessed for GO term enrichment with biological pathways using MSigDB. REViGO^89^ tree plots were generated in R using GO terms obtained from MSigDB.

### Weighted Gene Co-Expression Network Analysis

The CQN normalized expression values from bulk microglia were input to R WGCNA^31^ package v1.69. This analysis included 14,149 expressed genes, i.e. median(CQN) > 2. Modules were identified, their eigengenes were calculated and merged if correlation of eigengenes > 0.7. Genes in the 40 modules identified were tested for GO term enrichment via WGCNA. Module membership (MM) for each gene was calculated as the correlation between expression of each gene and its module eigengene. Genes with MM ≥ 0.7 are considered the hub genes for the network. Gene co-expression network plots were generated in Cytoscape v3.8 (http://www.cytoscape.org/). Each module eigengene was tested for association with age, sex and *APOE* ε4 carrier status independently using Pearson correlation. Co-expression network genes were annotated if they were significantly associated (p < 0.05) with the tested trait.

### Over-Representation and Correlation Analyses

Hypergeometric testing was performed in R to determine the enrichment of a select set of genes in previously reported signatures, bulk tissue expressed genes, WGCNA modules or 10X single cell clusters. Correlation between bulk tissue and bulk microglial normalized CQN data was calculated using Spearman’s rank correlation. Concordant and discordantly correlated genes were determined using the upper and lower quartiles from each dataset.

### Single Cell Data Analysis

For single cell RNA samples, 10X Genomics Cell Ranger Single Cell Software Suite v3.1.0^90^ was used to demultiplex raw base call files generated from the sequencer into FASTQ files. Raw reads were aligned to human genome build GRCh38. Reads aligned to gene transcript locus were counted to generate raw UMI counts per gene per barcode for each sample. The raw UMI matrices were filtered to only keep barcodes with > 500 UMIs and those that were classified as cells by Cell Ranger’s cell calling algorithm.

Quality control, normalization, clustering and marker gene identification were performed with Seurat v3^91^, followed by annotation of clusters using established cell type markers. We kept 1) barcodes with > 10% of UMI mapped to mitochondrial genome; 2) barcodes with < 400 or > 8000 detected genes; 3) barcodes with < 500 or > 46,425 mapped UMIs; 4) genes that are detected in < 5 cells. These thresholds were determined by UMI or gene distribution to identify undetectable genes and outlier barcodes that may encode background, damaged or multiple cells. UMI counts of remaining cells and genes were normalized using NormalizeData function, which gave natural log transformed expression adjusted for total UMI counts in each cell. The top 2000 genes whose normalized expression varied the most across cells were identified through FindVariableFeatures function with default parameters. Using those genes, cells from 6 samples were integrated using functions FindIntegrationAnchors and IntegrateData with default parameters. Principal components (PCs) of the integrated and scaled data were computed; and the first 31 PCs, which accounted for > 95% variance, were used in clustering cells. Cell clustering was performed using FindNeighbors and FindClusters with default parameters. Marker genes were identified in each cluster using FindMarkers in Seurat. Marker genes on one cluster must 1) be present in > 20% cells in the cluster; 2) the log(fold change) between expression in the cluster and other clusters must be > 0.25; 3) the rank sum test p-value (Bonferroni-adjusted) between cells in the cluster and cells in other clusters < 0.05.

### Meta-Analysis of Bulk Microglia RNA-seq Datasets

For these analyses we identified and obtained RNAseq data from three bulk microglial datasets that were available for download^16–18^ (**Supplementary Table S16**). We obtained sets of co-expression genes i.e. WGCNA modules from our own data for those modules the eigengenes of which were significantly correlated with age/sex/*APOE*. To determine whether these correlations replicated in the external datasets, we downloaded raw reads in FASTQ format for the three external bulk microglia datasets listed in **Supplementary Table S16**. Reads were mapped to human genome build hg38 and were counted and normalized in the same fashion as described previously. For each WGCNA module, we identified the central genes (i.e. genes whose Pearson correlation with module eigengene > 0.75). Using these central genes, module eigengenes were calculated in all datasets – ours and three external ones. We correlated module eigengenes with traits. Finally, we performed meta-analysis to combine correlations from multiple datasets using metacor function (random effect model) in R meta package.

### Joint Analysis of Single Cell RNA-seq Datasets

Raw single cell RNA-seq data from Olah et al (2020)^9^ was downloaded through Synapse (syn21438358) and processed through Cell Ranger. Quality control was performed as described prior to integration with our samples using Seurat v4.0.4. Integration was performed to combine the datasets by individual rather than sample. A total of 9 samples and 15,819 cells were retained from the Olah data^9^. Cells were annotated to include dataset of origin. Hypergeometric tests were performed to determine the enrichment of our microglial signature within each integrated cluster. Additionally, we also looked at overlap between integrated cluster genes and marker genes from our original single cell dataset. To identify where our original cells localized within the integrated dataset, we mapped the original cell IDs to the integrated clusters and calculated percentage overlap.

## Abbreviations

AD: Alzheimer’s disease
APOE: Apolipoprotein E
BM: Brodmann’s area
BSA: Bovine serum albumin
CERAD: Consortium to Establish a Registry for Alzheimer’s Disease
CNS: Central nervous system
CQN: Conditional quantile normalization
DAM: Disease-associated microglia
DPBS: Dulbecco’s phosphate buffered saline
FACS: Fluorescence-activated cell sorting
FPKM: Fragments per kilobase of transcript per million mapped reads
GEM: Gel bead-in emulsion
GO: Gene ontology
MACS: Magnetic-activated cell sorting
ME: Module eigengenes
MM: Module membership
PBS: Phosphate buffered saline
PC: Principal component
PCA: Principal component analysis
PFA: Paraformaldehyde
QC: Quality control
RNAseq: RNA sequencing
ROSMAP: Rush University Religious Order Study-Memory and Aging Project
scRNAseq: Single cell RNA sequencing
snRNAseq: Single nuclei RNA sequencing
UMI: Unique molecular identifier
WGCNA: Weighted gene co-expression network analysis

## Acknowledgements

The authors thank the patients and their families for their participation, without whom these studies would not have been possible.

This study was supported by NIH funding U01 AG046193, RF1 AG051504, R01 AG051504 to NET.

We thank our colleagues in the neurosurgery team Christopher Louie, Karim ReFaey and Ivan Segura Duran. We thank our colleagues at the Mayo Clinic Genome Analysis Core (GAC) for their collaboration, particular gratitude to Bruce Eckloff and Julie Lau. We also acknowledge the AMP-AD RNAseq reprocessing team, in particular Dr. Kirsten Dang, Dr. Thanneer Perumal and Dr. Ben Logsdon at Sage Bionetworks.

**AMP-AD RNASeq datasets:** This study is a cross-consortia project using RNAseq data generated through grants U01AG046152, U01AG046170, and U01AG046139. **For the Mayo RNAseq study:** The results published here are in whole or in part based on data obtained from the AD Knowledge Portal (https://adknowledgeportal.synapse.org/). Study data were provided by the following sources: The Mayo Clinic Alzheimers Disease Genetic Studies, led by Dr. Nilüfer Ertekin-Taner and Dr. Steven G. Younkin, Mayo Clinic, Jacksonville, FL using samples from the Mayo Clinic Study of Aging, the Mayo Clinic Alzheimers Disease Research Center, and the Mayo Clinic Brain Bank. Data collection was supported through funding by NIA grants P50 AG016574, R01 AG032990, U01 AG046139, R01 AG018023, U01 AG006576, U01 AG006786, R01 AG025711, R01 AG017216, R01 AG003949, NINDS grant R01 NS080820, CurePSP Foundation, and support from Mayo Foundation. Study data includes samples collected through the Sun Health Research Institute Brain and Body Donation Program of Sun City, Arizona. The Brain and Body Donation Program is supported by the National Institute of Neurological Disorders and Stroke (U24 NS072026 National Brain and Tissue Resource for Parkinson’s Disease and Related Disorders), the National Institute on Aging (P30 AG19610 Arizona Alzheimer’s Disease Core Center), the Arizona Department of Health Services (contract 211002, Arizona Alzheimer’s Research Center), the Arizona Biomedical Research Commission (contracts 4001, 0011, 05-901 and 1001 to the Arizona Parkinson’s Disease Consortium) and the Michael J. Fox Foundation for Parkinson’s Research. **For the ROSMAP study**: The results published here are in whole or in part based on data obtained from the AD Knowledge Portal (https://adknowledgeportal.synapse.org).

Study data were provided by the Rush Alzheimer’s Disease Center, Rush University Medical Center, Chicago. Data collection was supported through funding by NIA grants P30AG10161 (ROS), R01AG15819 (ROSMAP; genomics and RNAseq), R01AG17917 (MAP), R01AG30146, R01AG36042 (5hC methylation, ATACseq), RC2AG036547 (H3K9Ac), R01AG36836 (RNAseq), R01AG48015 (monocyte RNAseq) RF1AG57473 (single nucleus RNAseq), U01AG32984 (genomic and whole exome sequencing), U01AG46152 (ROSMAP AMP-AD, targeted proteomics), U01AG46161(TMT proteomics), U01AG61356 (whole genome sequencing, targeted proteomics, ROSMAP AMP-AD), the Illinois Department of Public Health (ROSMAP), and the Translational Genomics Research Institute (genomic). Additional phenotypic data can be requested at www.radc.rush.edu. **For the MSBB study:** The results published here are in whole or in part based on data obtained from the AD Knowledge Portal (https://adknowledgeportal.synapse.org/). These data were generated from postmortem brain tissue collected through the Mount Sinai VA Medical Center Brain Bank and were provided by Dr. Eric Schadt from Mount Sinai School of Medicine.

## Data Sharing Statement

The data in this manuscript are available via the AD Knowledge Portal (https://adknowledgeportal.synapse.org). The AD Knowledge Portal is a platform for accessing data, analyses and tools generated by the Accelerating Medicines Partnership (AMP-AD) Target Discovery Program and other National Institute on Aging (NIA)-supported programs to enable open-science practices and accelerate translational learning. The data, analyses and tools are shared early in the research cycle without a publication embargo on secondary use. Data is available for general research use according to the following requirements for data access and data attribution (https://adknowledgeportal.synapse.org/DataAccess/Instructions).

| Dataset | Data Type | Description | SynapseID | DoD |
| --- | --- | --- | --- | --- |
| Mayo RNAseq TCX | Metadata | Individual human and RNAseq | syn5550404 | na |
| Mayo RNAseq CER | Metadata | Individual human and RNAseq | syn5550404 | na |
| Mayo RNAseq TCX | RNASeq Expression | Consensus processed RNASeq raw counts | syn8690799 | 10/2/2019 |
| Mayo RNAseq CER | RNASeq Expression | Consensus processed RNASeq raw counts | syn8690904 | 10/2/2019 |
| ROSMAP | Metadata | ID Key | syn3382527 | 10/2/2019 |
| ROSMAP | Metadata | Individual human | syn3191087 | 10/2/2019 |
| ROSMAP | Metadata | Assay RNAseq | syn21088596 | 1/2/2020 |
| ROSMAP | RNASeq Expression | Consensus processed RNASeq raw counts | syn8691134 | 10/2/2019 |
| MSBB | Metadata | Individual human | syn6101474 | 11/22/2019 |
| MSBB | Metadata | Assay RNAseq | syn6100548 | 10/2/2019 |
| MSBB | RNASeq Expression | Consensus processed RNASeq raw counts | syn8691099 | 10/2/2019 |

**Data from AD knowledge portal utilized in this study.** DoD = Date of download, “na” indicates data that was generated by study authors and shared within the AD knowledge portal. Data were obtained from the RNAseq Harmonization Study in the AD knowledge portal (DOI: https://doi.org/10.7303/syn9702085).

## Author Contributions

TP and NET wrote the manuscript; NET and MA designed the study; TP, XW and ZQ performed data analysis; JC consulted on statistical methods; TP, TPC, XW, YM, RMA generated tables and figures; EM, CAG, SG, KC, RW, HGC and AQH provided neurosurgical tissue samples; TP, TPC, LJLT, SJL, SL, FQTN, CCGH, KGM, and TN performed experimental procedures from blood and tissue samples. All authors read the manuscript and provided input and consultation. NET oversaw the study and provided direction, funding and resources.

