## Supplemental Figures for "Transcriptional landscape of human microglia reveals robust gene expression signatures that implicates age, sex and *APOE*-related immunometabolic pathway perturbations"

**Number of Figures:** 4

**Number of Supplementary Tables**: 20

**Number of Supplementary Figures**: 10

**Word Count**: 5,179 (excludes Methods word count of 2,155)

**References**: 91

**Supplementary Figures**

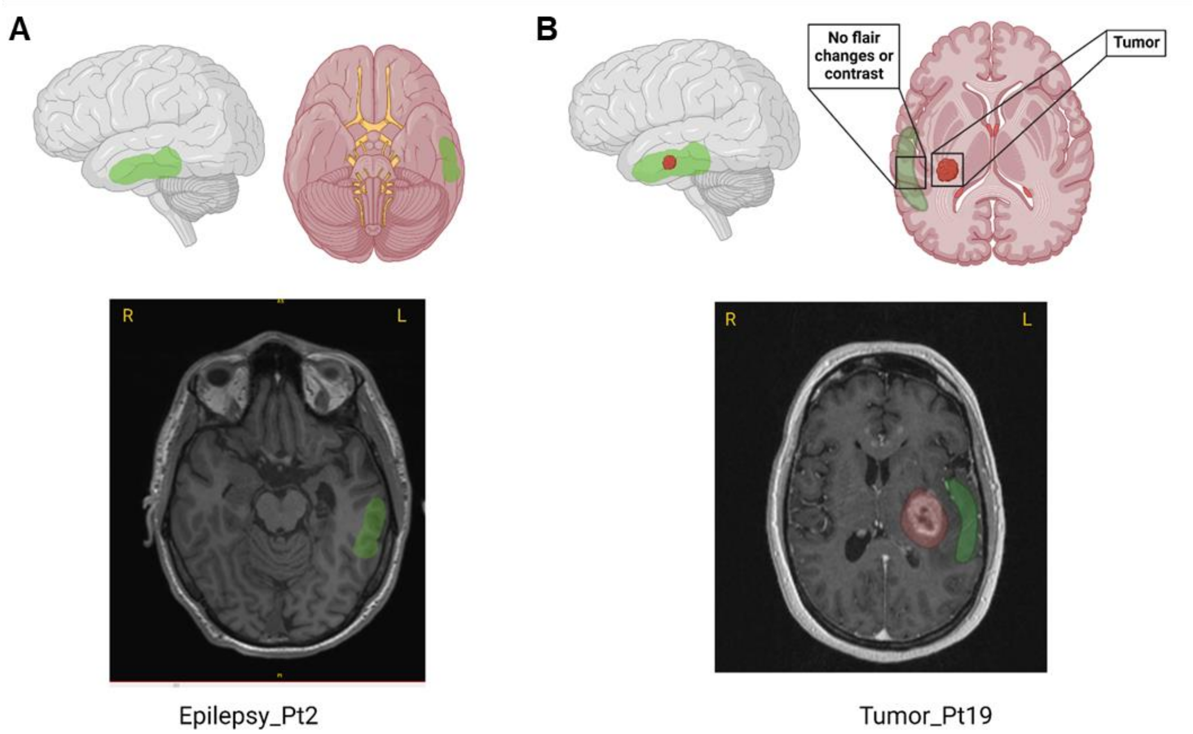


**Figure S1. Surgical approaches for excision of non-affected brain tissue.** (A) 3-D model and axial T1 MRI image highlighting general areas where normal tissue was obtained in epilepsy cases (green). (B) 3-D model and axial T1 MRI image highlighting the area for normal tissue removal in patients with oncologic resections (green) [Created with BioRender.com].

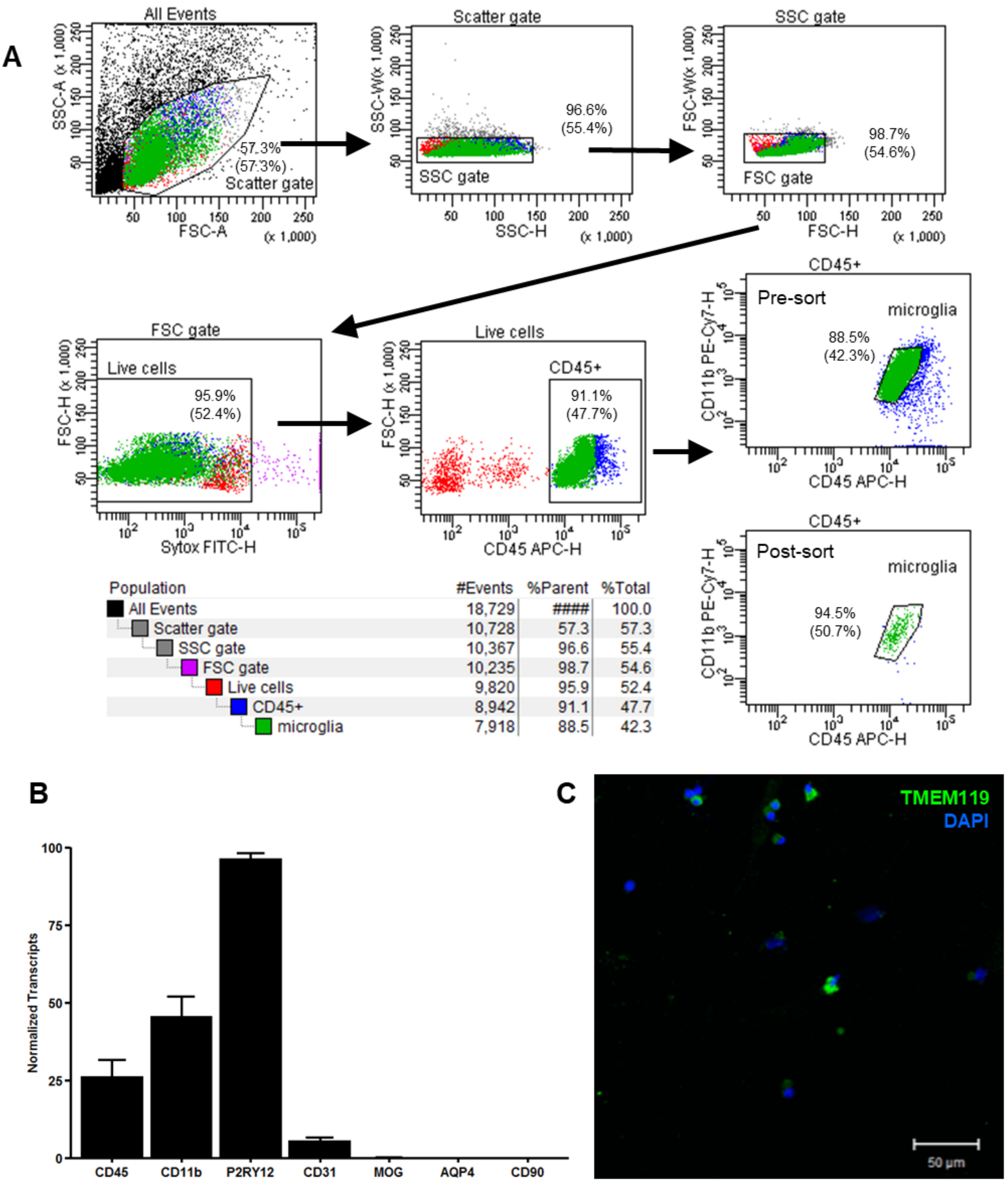


**Figure S2. Validation of sorted cells indicate enriched microglial population.** (A) Dot plots showing FACS gating strategy for isolating $CD11b^{+}/CD45^{intermediate}$ cell population. Sorted cells underwent bulk and single cell RNA sequencing. (B) Aggregated qPCR of normalized transcripts from sorted cells for 19 bulk microglia samples. These cells express the expected microglial signature and are $P2RY12^{+}$. Other cell type markers are not expressed. (C) Immunocytochemistry image of sorted cells stained with anti-TMEM119 conjugated to Goat anti-rabbit Alexa 488 (green) and DAPI (blue). Double stained cells represent $TMEM119^{+}$ microglia (83% of total cells).

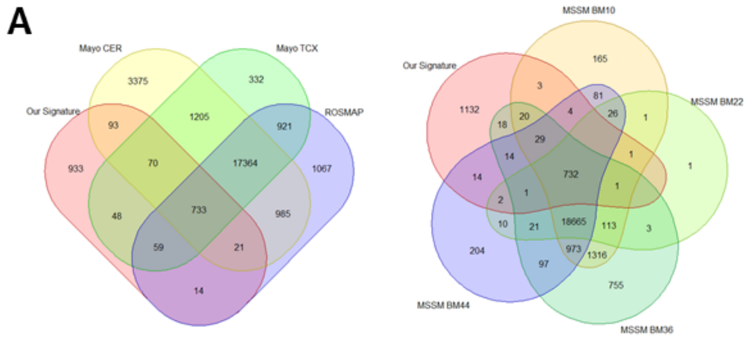


**B**

| Dataset | # Samples | # Expressed Genes | % Overlap | Region |
|---|---|---|---|---|
| Our Signature | 19 | 1971 | | |
| Mayo CER | 144 | 20732 | 4.4% | Cerebellum |
| Mayo TCX | 148 | 23846 | 3.8% | Temporal cortex |
| MSSM BM10 | 160 | 22130 | 3.6% | Frontal pole |
| MSSM BM22 | 133 | 19577 | 3.8% | Superior temporal gyrus |
| MSSM BM36 | 119 | 22758 | 3.6% | Parahippocampal gyrus |
| MSSM BM44 | 133 | 20873 | 3.8% | Inferior frontal gyrus |
| ROSMAP DLPFC | 469 | 21164 | 3.9% | Dorsolateral prefrontal cortex |

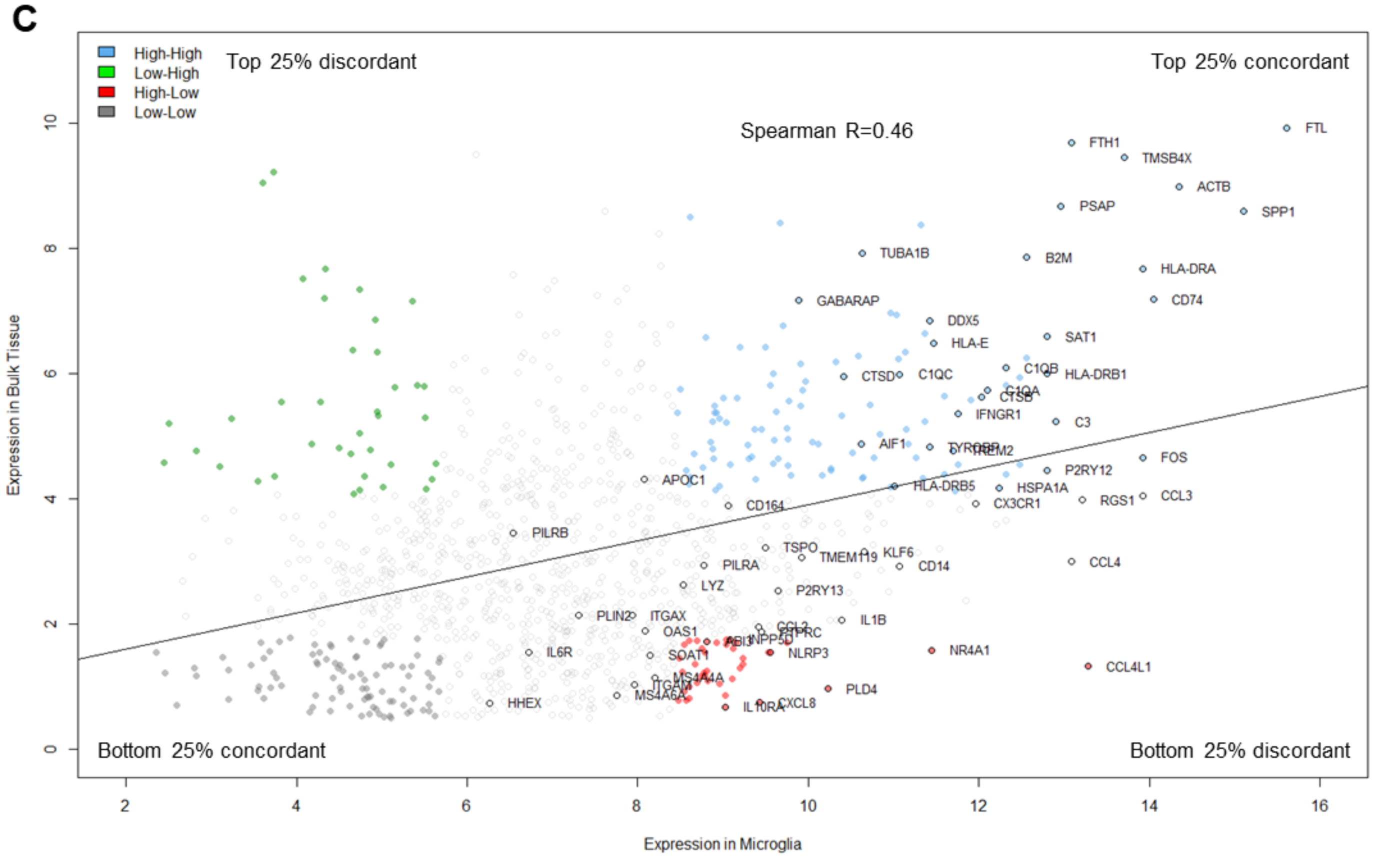


**Figure S3. Correlation of our microglial signature vs bulk surgical tissue.** (A) Venn diagrams showing the number of overlapping genes between our microglial signature and the 7 AMP-AD brain regions. (B) Total number of genes expressed per region and % bulk genes representing the microglial signature (overlap). (C) Scatter plot showing correlation of expression in CQN values for our microglial signature genes in bulk microglia vs. bulk surgical fresh brain tissue. Concordant and discordant genes in the highest and lowest expression quadrants are highlighted in blue (high-high), green (low-high), red (high-low) and
22 gray (low-low). Select genes are annotated (o).

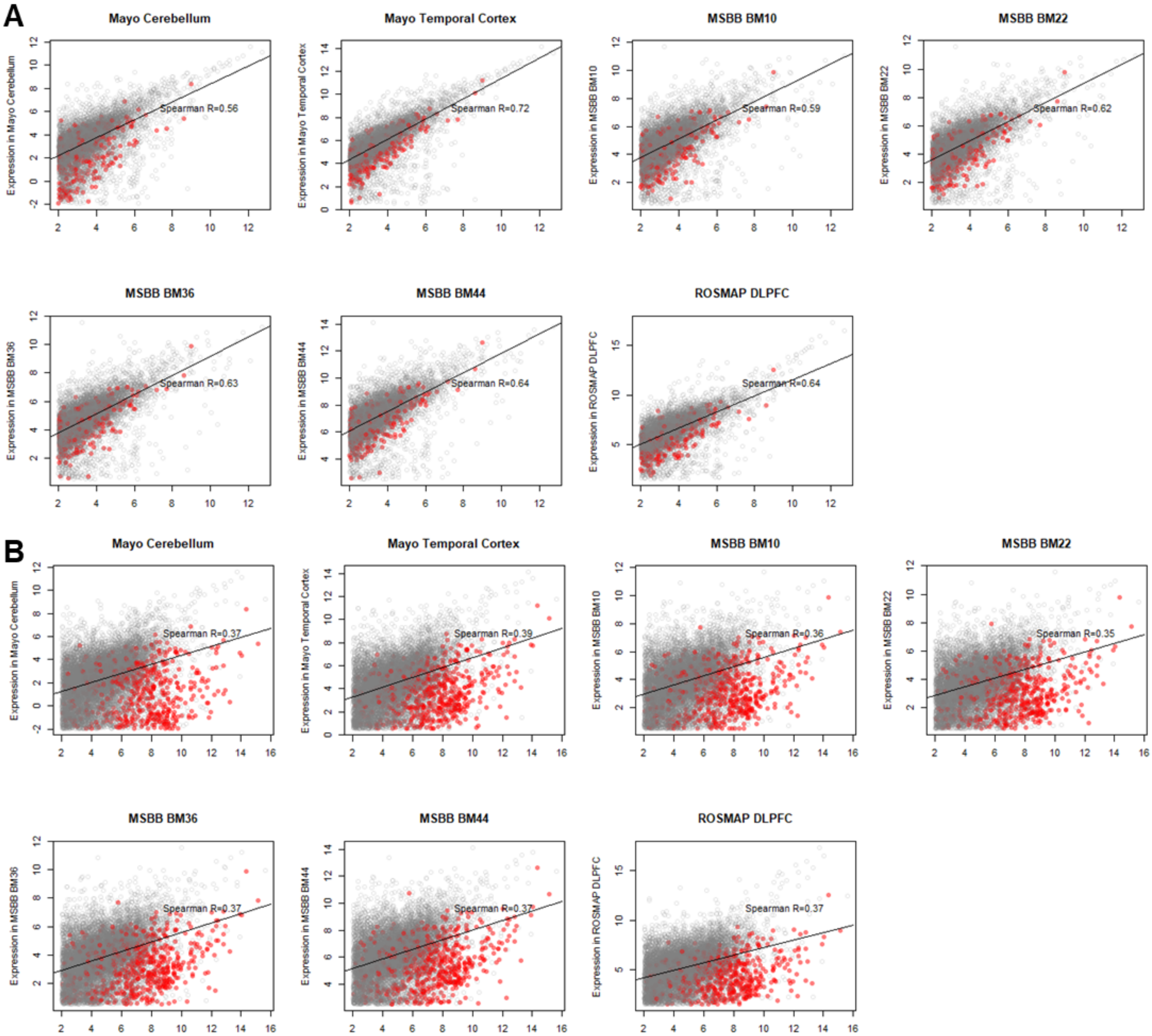


**Figure S4. Correlation of AMP-AD bulk tissue transcriptomic datasets with bulk surgical and microglial data.** (A) Scatter plots showing Spearman's correlation of gene expression between our bulk surgical tissue data and the 7 AMP-AD bulk tissue RNA-seq datasets using CQN. (B) Scatter plots showing Spearman's correlation of our bulk microglial data and the AMP-AD bulk tissue datasets using CQN. BRETIGEA microglial marker genes are highlighted in red.

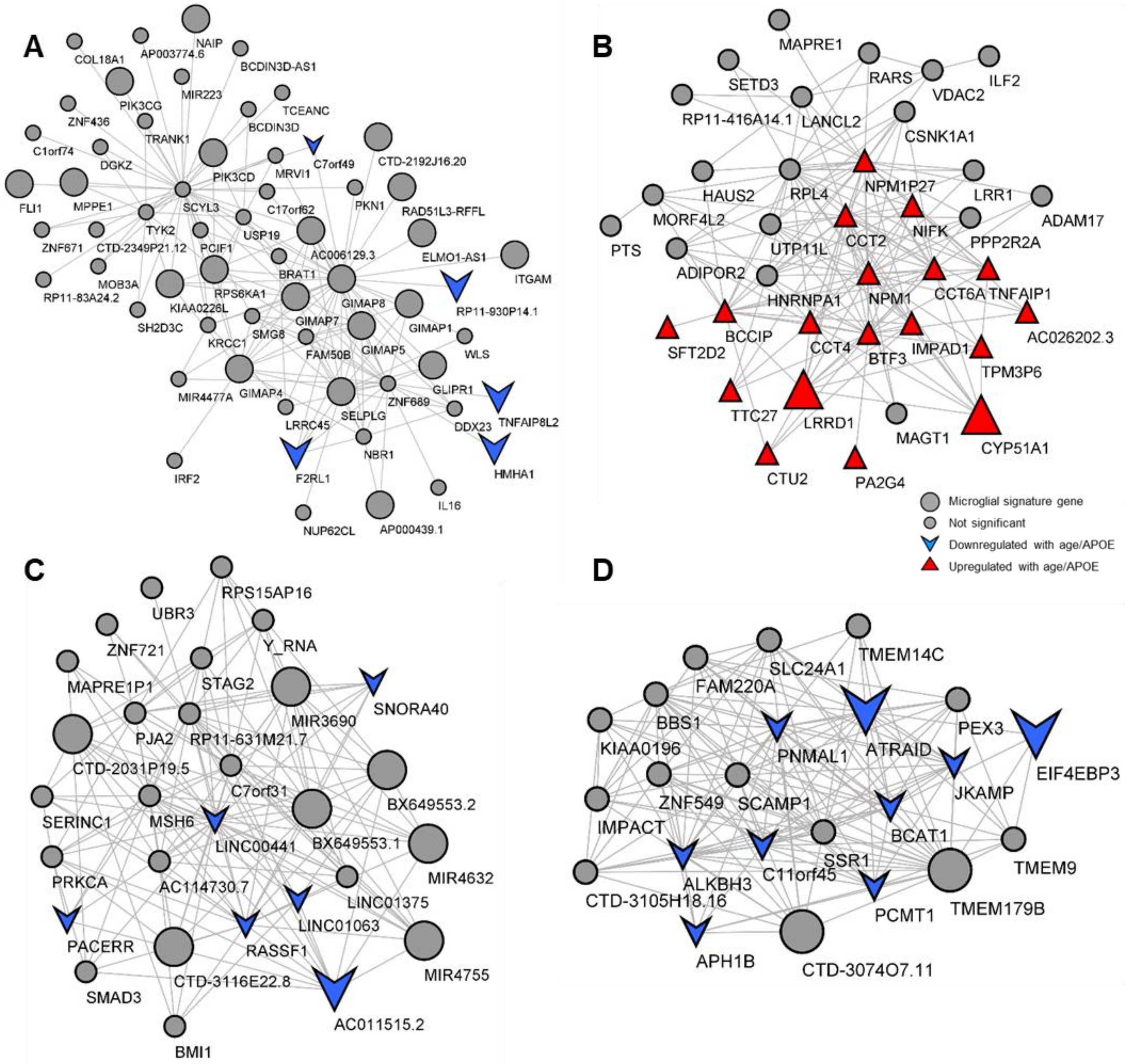


**Figure S5. Co-expression networks of WGCNA modules significantly associated with age, sex or *APOE*.** Genes with module membership > 0.7 are shown here with microglial signature genes denoted with larger circles, upregulated genes with age or *APOE* shown as red triangles and downregulated genes shown as as blue inverted arrows. (A) Module 4 co-expression network which was significantly downregulated in *APOE* ε4 carriers. (B) Module 28 gene co-expression network was upregulated in *APOE* ε4 carriers. (C) Module 34 gene co-expression network was downregulated in *APOE* ε4 carriers. (D) Module 36 gene co-expression network was downregulated in *APOE* ε4 carriers.

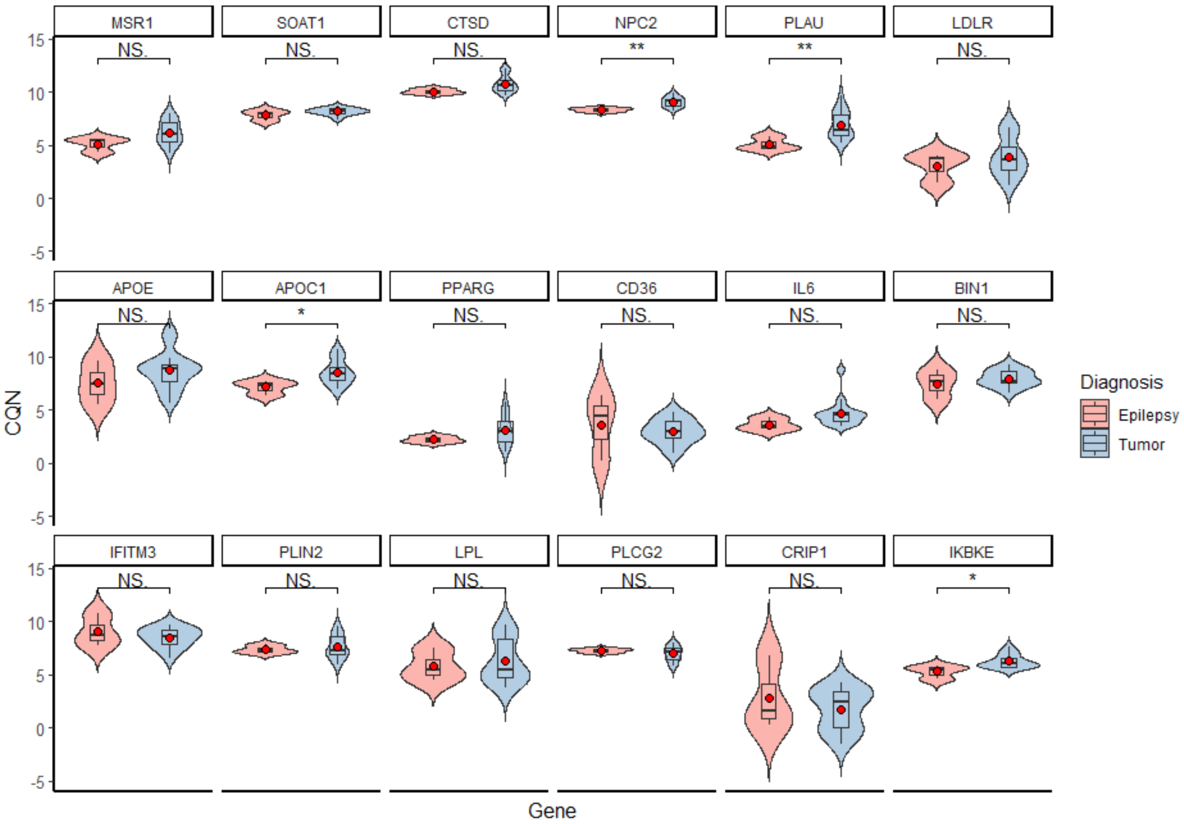


**Figure S6. Expression levels of genes of interest stratified by diagnosis.** Violin plots showing the expression of key genes within tumor and epilepsy samples demonstrate that most genes are not affected by diagnosis.

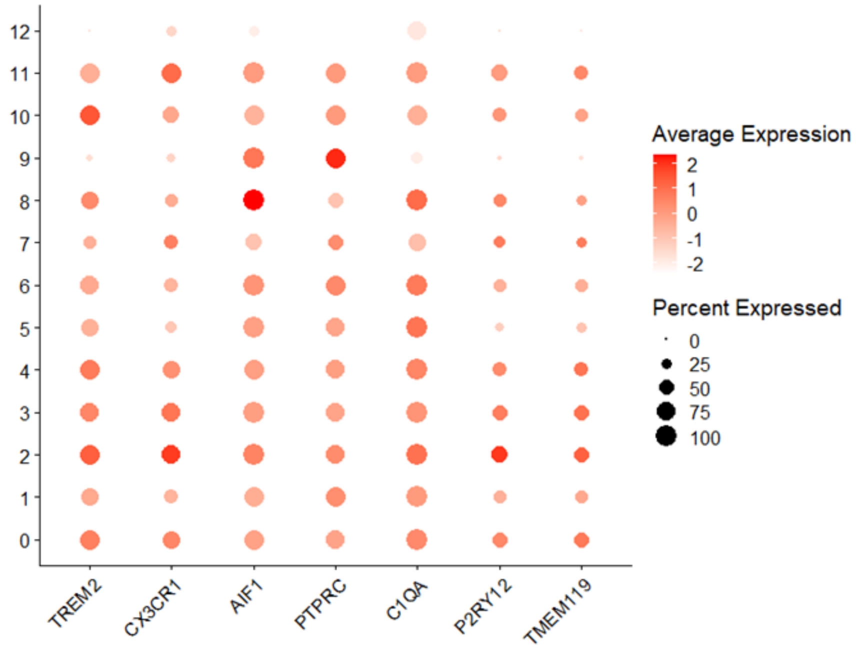


**Figure S7. Expression of established microglial marker genes in single cell data.** Dot plot showing expression of established microglial marker genes from the literature across all clusters.

26

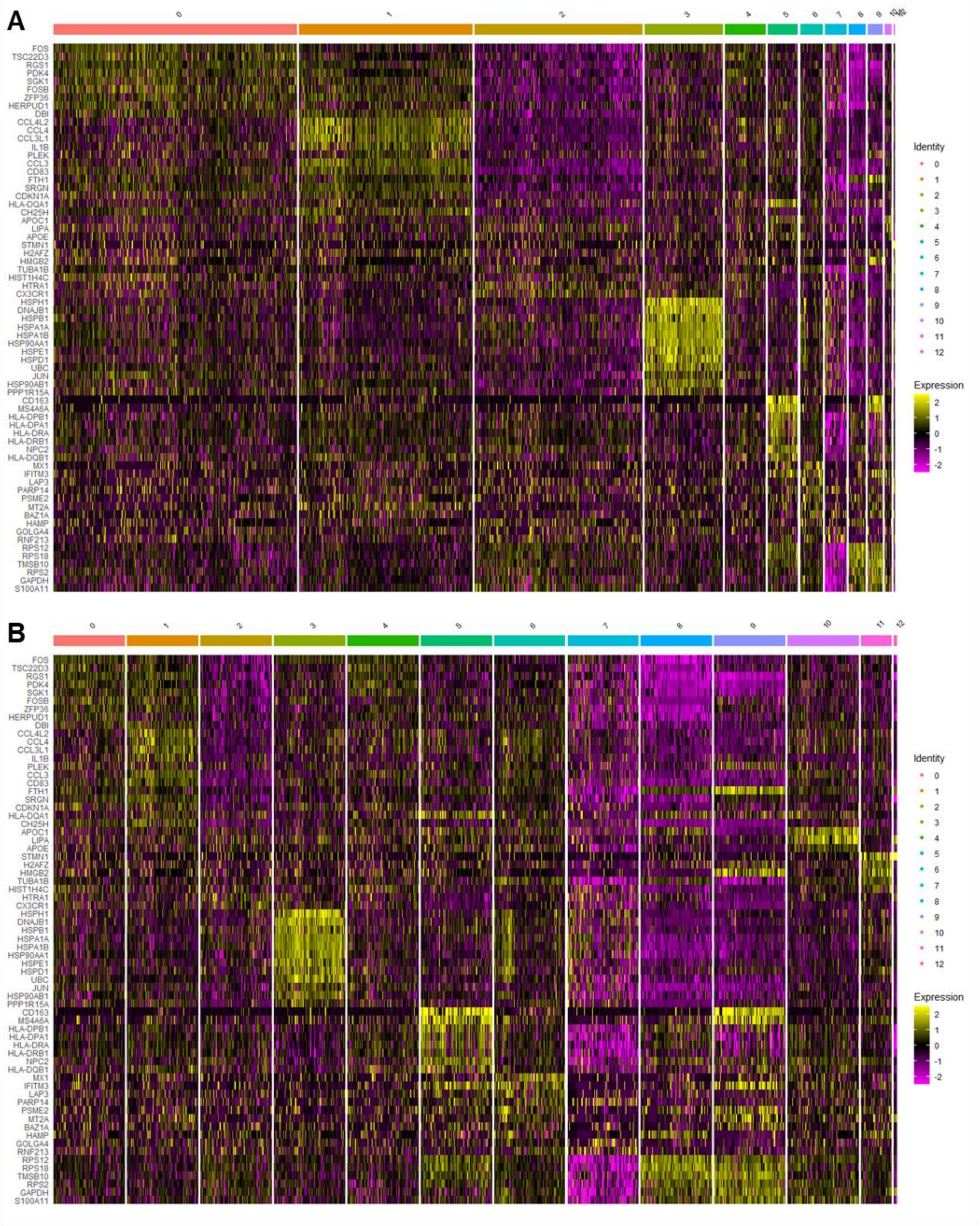


**Figure S8. Expression of top cluster marker genes across single cell clusters.** (A) Heatmap showing expression of selected marker genes (log fold change < -0.1 or > 0.1; q<0.05) for myeloid clusters. (B) Same heatmap downsampled to 100 cells.

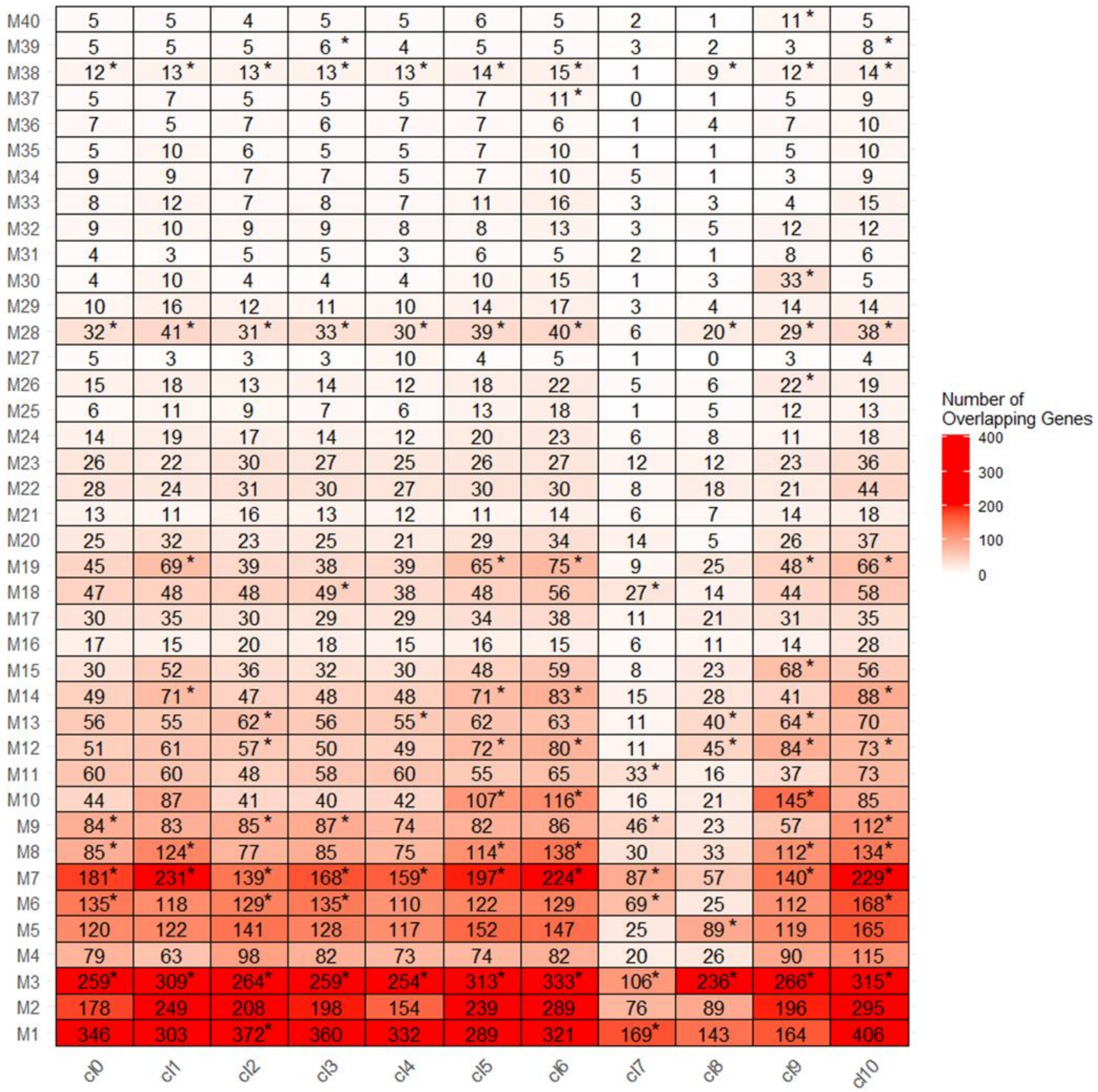


**Figure S9. Hypergeometric overlap of module genes across all single cell clusters.** Hypergeometric distribution of enrichment between module genes and clusters showing numbers of overlapping genes. * represents module genes that were significantly enriched in the cluster ($p < 0.05$).

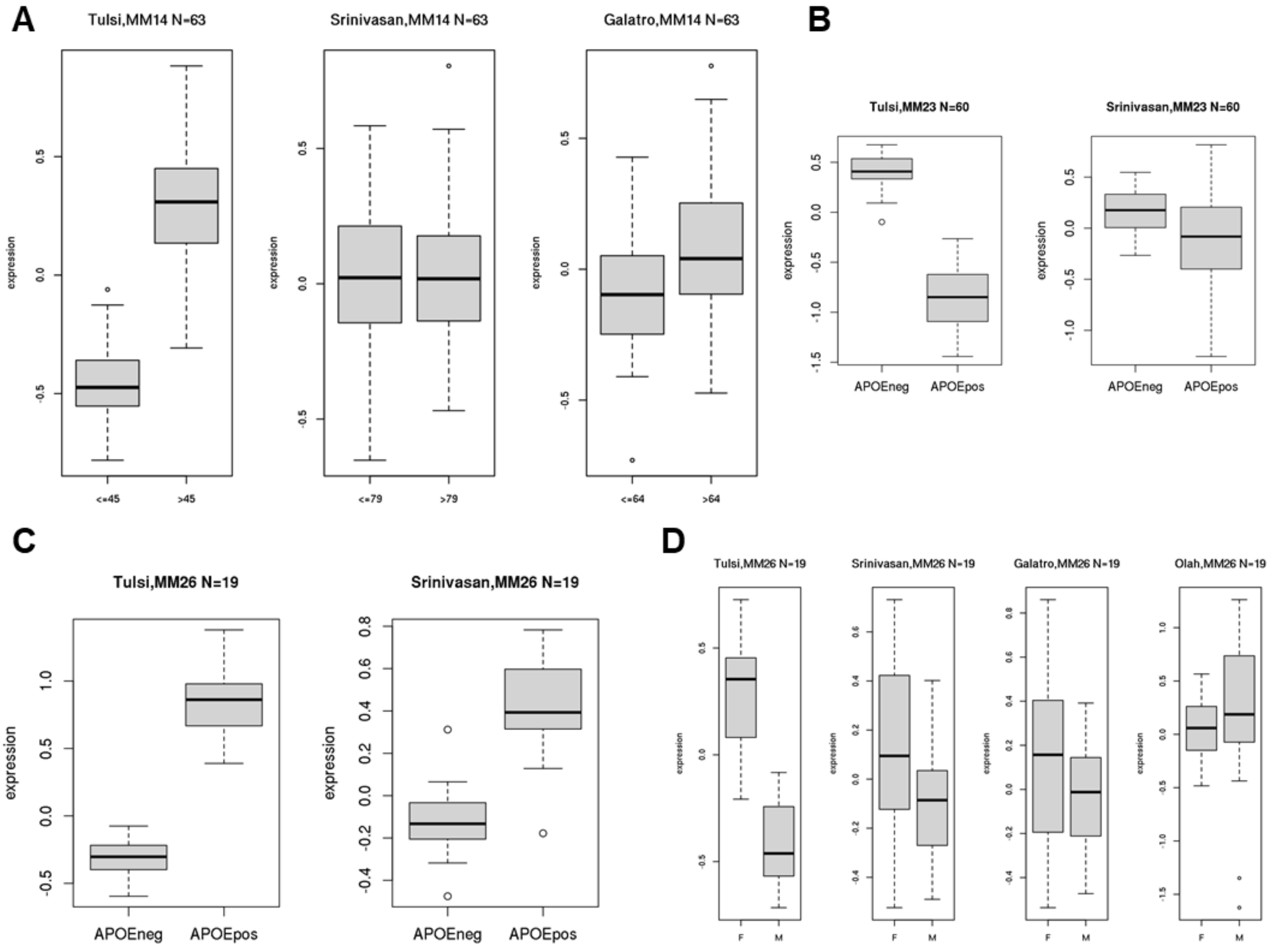


**Figure S10. WGCNA meta-analysis showing all expression of all genes within modules.** Box plots show the expression patterns between traits for all genes filtered by module membership (MM>0.75) within the modules of interest. (A) Module 14 genes for datasets stratified by median age. (B) Module 23 genes stratified by *APOE* ε4 carrier status shows similar trends between datasets. (C) Module 26 genes show similar expression changes for *APOE* ε4 carriers and non-carriers between datasets. (D) Module 26 genes show similar gene expression trends for most datasets except Olah et al (2018).

N= Number of genes plotted
